# Subpopulations of neurons in lOFC encode previous and current rewards at time of choice

**DOI:** 10.1101/2021.05.06.442972

**Authors:** David Hocker, Carlos D. Brody, Cristina Savin, Christine M Constantinople

## Abstract

Studies of neural dynamics in lateral orbitofrontal cortex (lOFC) have shown that subsets of neurons that encode distinct aspects of behavior, such as value, may project to common downstream targets. However, it is unclear whether reward history, which may subserve lOFC’s well-documented role in learning, is represented by functional subpopulations in lOFC. Previously, we analyzed neural recordings from rats performing a value-based decision-making task, and we documented trial-by-trial learning that required lOFC (Constantinople *et al*., 2019). Here we characterize functional subpopulations of lOFC neurons during behavior, including their encoding of task variables. We found five distinct clusters of lOFC neurons, either based on clustering of their trial-averaged peristimulus time histograms (PSTHs), or a feature space defined by their average conditional firing rates aligned to different task variables. We observed weak encoding of reward attributes, but stronger encoding of reward history, the animal’s left or right choice, and reward receipt across all clusters. Only one cluster, however, encoded the animal’s reward history at the time shortly preceding the choice, suggesting a possible role in integrating previous and current trial outcomes at the time of choice. This cluster also exhibits qualitatively similar responses to identified corticostriatal projection neurons in a recent study (Hirokawa *et al*., 2019), and suggests a possible role for subpopulations of lOFC neurons in mediating trial-by-trial learning.

## 2 Introduction

Previous experience can profoundly influence subsequent behavior and choices. In trial-based tasks, the effects of previous choices and outcomes on subsequent ones are referred to as “sequential effects,” and while they are advantageous in dynamic environments, they produce suboptimal biases when outcomes on each trial are independent. The orbitofrontal cortex (OFC) has been implicated in updating behavior based on previous experience, particularly when task contingencies are partially observable [1–4]. However, it is unclear whether behavioral flexibility in OFC is mediated by dedicated subpopulations of neurons exhibiting distinct encoding and/or connectivity. We previously trained rats on a value-based decision-making task, in which they chose between explicitly cued, guaranteed and probabilistic rewards on each trial [5, 6]. Despite the fact that outcomes were independent on each trial, we observed several distinct sequential effects that contributed to behavioral variability. Optogenetic perturbations of the lateral orbitofrontal cortex (lOFC) eliminated one particular sequential effect, an increased willingness to take risks following risky wins, but spared other types of trial-by-trial learning, such as spatial “win-stay/lose-switch” biases. We interpreted this data as evidence that (1) different sequential effects may be mediated by distinct neural circuits, and (2) lOFC promotes learning of abstract biases that reflect the task structure (here, biases for the risky option), but not spatial ones [6].

Electrophysiological recordings from lOFC during this task revealed encoding of reward history and reward outcomes on each trial at the population level, which could in principle support sequential effects [6]. Recent studies of rodent OFC have suggested that despite the apparent heterogeneity of neural responses in prefrontal cortex, neurons can be grouped into distinct clusters that exhibit similar task-related responses and, in some cases, project to a common downstream target [7, 8]. In light of these results, we hypothesized that reward history might be encoded by a distinct cluster of neurons in lOFC. This would suggest that the lOFC-dependent sequential effect we observed (an increased willingness to take risks following risky wins) may derive from a subpopulation of neurons encoding reward history, and may potentially project to a common downstream target.

Here, we analyzed an electrophysiological dataset from lOFC during a task in which independent and variable sensory cues conveyed dissociable reward attributes on each trial (reward probability and amount; [5, 6]). Our previous analysis investigated single units responses for sensitivity to task parameters, and here we found that clustering of lOFC neurons based on either their trial-averaged peristimulus time histograms (PSTHs), or a feature space defined by their average conditional firing rates aligned to different task variables [7], both revealed five clusters of neurons with different response profiles. We exploited the temporal variability of task events to fit a generalized linear model (GLM) to lOFC firing rates, generating a rich description of the encoding of various task variables over time in individual neurons. All of the clusters exhibited weak encoding of reward attributes, and stronger encoding of reward history, the animal’s left or right choice, and reward receipt. Encoding of these variables was broadly distributed across clusters. Only one cluster, however, encoded the animal’s reward history at the time shortly preceding the choice. This distinct encoding was observable by three independent metrics (coefficient of partial determination, mutual information, and discriminability or *d*′) and two separate clustering methods. Intriguingly, this cluster exhibited qualitatively similar responses to identified corticostriatal cells in a previous study [7], and also exhibited the strongest encoding of reward outcomes, suggesting that these neurons are well-situated to integrate previous and current reward experience. We hypothesize that this subpopulation of neurons, which were identifiable based on their temporal response profiles alone, may mediate sequential learning effects by integrating previous and current trial outcomes at the time of choice.

## 3 Results

Rats’ behavior on this task has been previously described [5, 6]. Briefly, rats initiated a trial by poking their nose in a central nose port (Fig. 1A-C). They were then presented with a series of pseudo-randomly timed light flashes, the number of which conveyed information about reward probability on each trial. Simultaneously, they were presented with randomly timed auditory clicks, the rate of which conveyed the volume of water reward (6-48 *μ*l) baited on each side. After a cue period of ~ 2.5 – 3s, rats reported their choice by poking in one of the two side ports. Reward volume and probability of reward (*p*) were randomly chosen on each trial. On each trial, the left or right port was randomly designated as risky (*p* < 1) or safe (*p* = 1). Well-trained rats tended to choose the side offering the greater subjective value, and trial history effects contributed to behavioral variability [5, 6]. We analyzed 659 well-isolated single-units in the lOFC with minimum firing rates of 1Hz, obtained from tetrode recordings [6].

**Figure 1:**
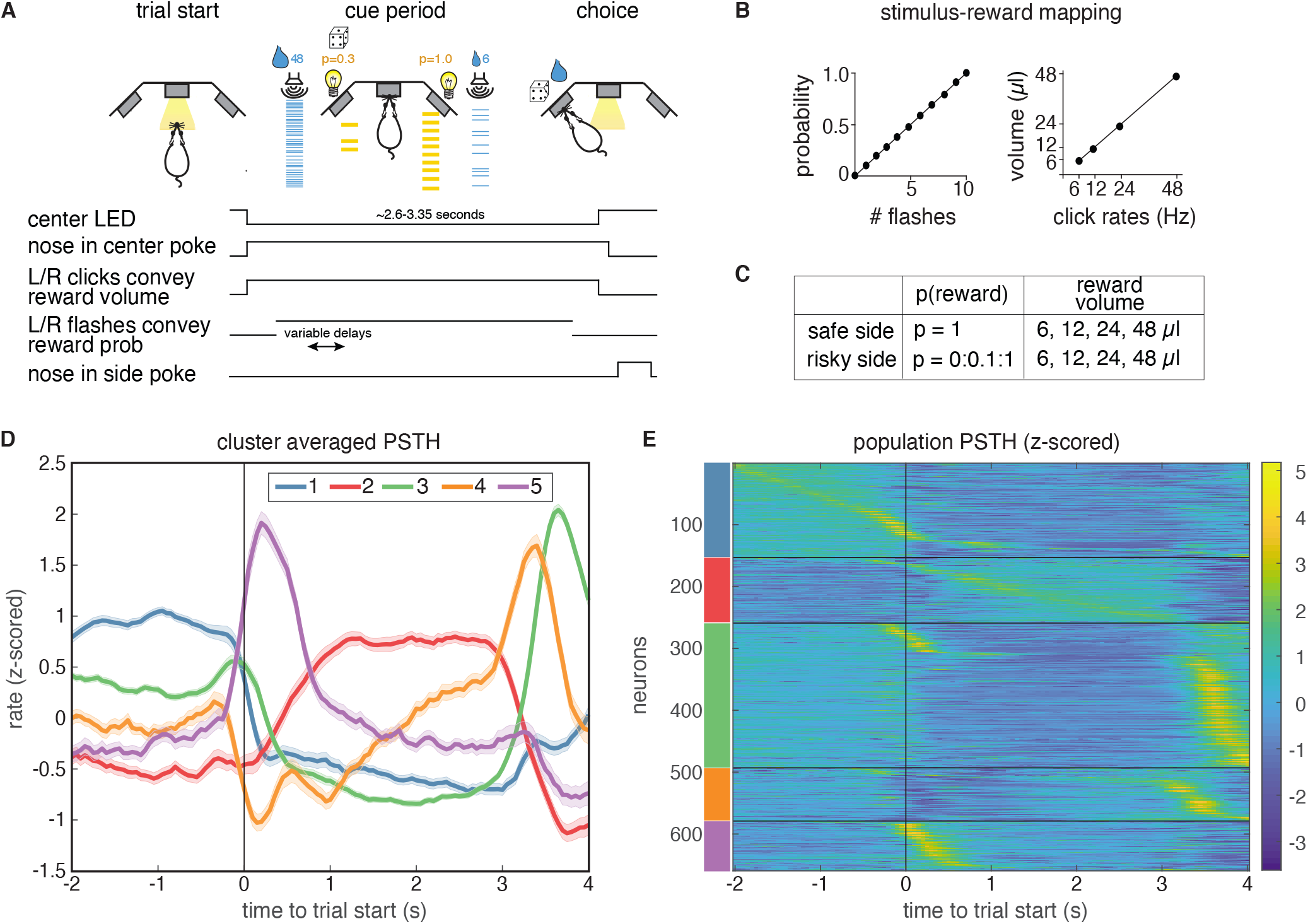
A. Behavioral task: rats chose between a guaranteed and probabilistic water reward on each trial, that could each be one of four water volumes. Rats were well-trained and tended to choose the option with the greater subjective value. Reward probability and volume were cued by visual flashes and auditory click rates, respectively. B. Mapping between the visual flashes/auditory clicks and reward attributes. C. Range of reward attributes. D. Cluster-specific, mean responses of the trial-averaged, z-scored PSTHs. Error bars denote s.e.m. E. Z-scored PSTHs for neurons in each cluster, sorted by time to peak within each cluster. Colored bars on the left indicate the cluster identities from panel D. Panels A-C reproduced and modified from [6].

### 3.1 Single units in lOFC belong to clusters with distinct temporal response profiles

To characterize the temporal dynamics of the neural responses during this task, we performed k-means clustering on trial-averaged PSTHs (“PSTH clustering”), aligned to trial initiation. We used the gap statistic [9], which quantifies the improvement in cluster quality with additional clusters relative to a null distribution (Methods), to choose a principled number of clusters, and identified five distinct clusters of responses (Fig. 1D). Each cluster has a stereotyped period of elevated activity during the task: at trial start, during the cue period, and at or near reward delivery, with the largest cluster (cluster 3) having activity just before the rat entered the reward port (Suppl. Fig. S1). Cluster 2 is the only cluster that exhibited persistent activity during the cue period. These data indicate that despite the well-documented heterogeneity of neural responses in prefrontal cortex, responses in lOFC belong to one of a relatively small, distinct set of temporal response profiles aligned to different task events.

A recent study in rat lOFC similarly found discrete clusters of neural responses by clustering the conditional firing rates of neurons for different task-related variables (“conditional clustering”) [7]. We sought to compare results from clustering PSTHs and conditional firing rates. Therefore, we generated conditional response profiles for each neuron (Fig. 2B; Methods), and performed clustering on these conditional responses via the same procedure. This also revealed five distinct clusters that exhibited qualitatively similar temporal response profiles as the clusters that were based on the average PSTHs (Fig. 2C-E, Suppl. Fig. S2). Moreover, clusters 2 and 3 were comprised of highly similar groups of neurons across both procedures, where similarity was quantified by the consistency of being labeled in a given cluster across clustering approaches (Fig. 2E, 66% overlap of cluster 2 neurons and 70% of cluster 3 neurons). This indicates that neurons in these two clusters are robustly identified by either their temporal response profiles or their encoding of task variables (Fig. 2E).

**Figure 2:**
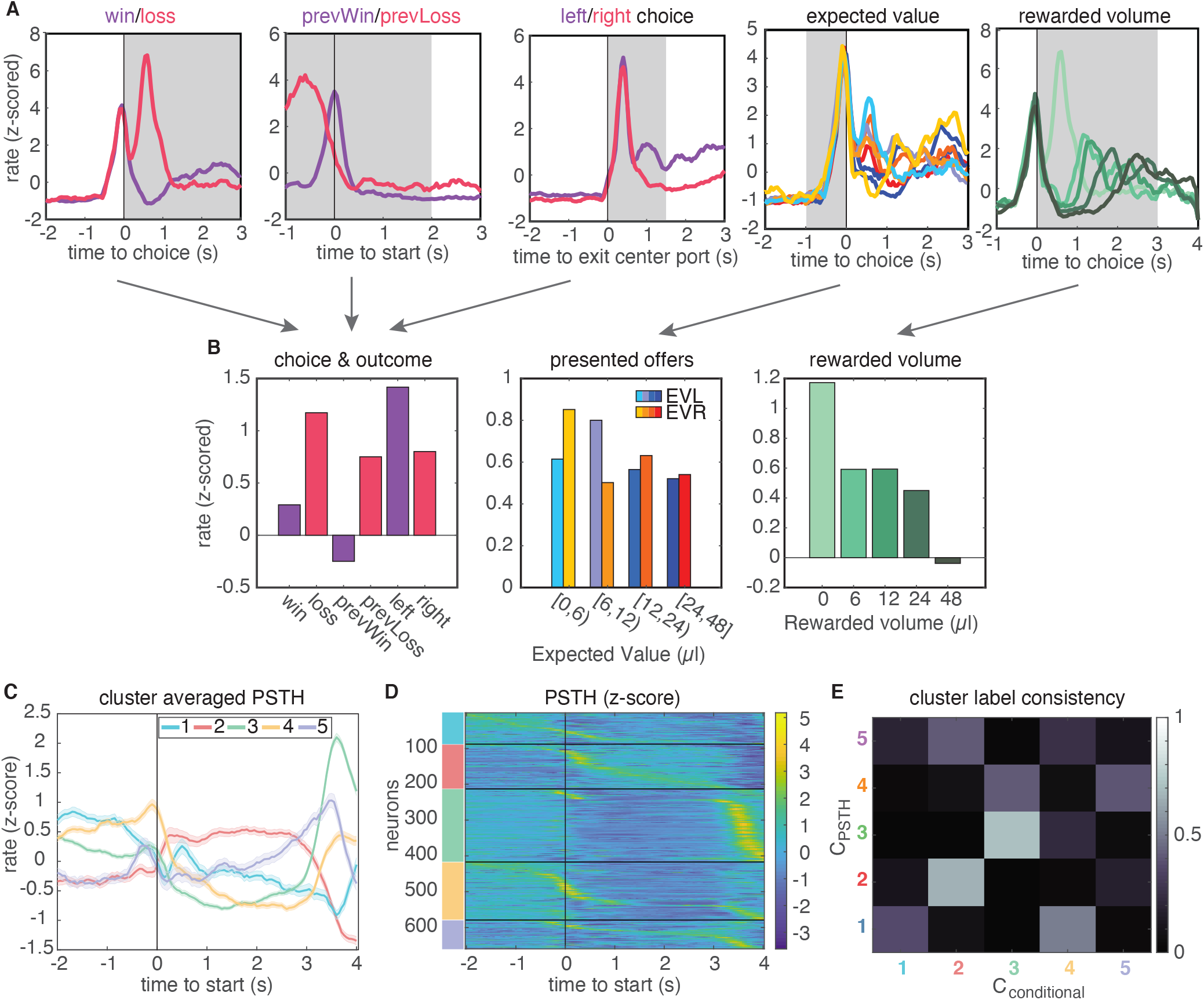
A. Feature-conditioned PSTHs of an example neuron. The average z-scored firing rate in each gray time window comprises an element in the feature space. B. Feature space for conditional clustering for the sample neuron from A. The average firing rate for each condition is concatenated to yield a 19-dimensional feature space. Here, features are grouped into three qualitative types (for presentation purposes only): responses for choice and trial outcome (left); responses to expected value of left and right offers (middle); and responses to received reward volume (right). C. Cluster-averaged PSTHs, aligned to trial initiation, using this conditional feature space. Error bars denote s.e.m. D. Z-scored PSTHs for all neurons, sorted by time to peak within each cluster. E. Consistency of cluster labeling, calculated as the conditional probability *P*(*C*_conditional_|*C*_PSTH_) of belonging in any ‘conditional’ cluster (*C*_conditional_), given that a unit belongs to a certain PSTH-defined cluster (*C*_PSTH_).

We evaluated the quality of clustering using two additional methods. First, we visualized clustering using t-SNE projections of data into two dimensions, and further quantified cluster separation using the Mahalanobis distance among clusters, which in this context can be interpreted as an average z-score for data in cluster from a given cluster mean (Suppl. Fig. S3). We found that lOFC responses obtained by both procedures were qualitatively clustered in feature space, and demonstrated Mahalahobis distances with smaller intracluster distances than across-cluster distances. Second, we used the PAIRS statistic to more directly test whether lOFC responses belong to distinct clusters [10]. PAIRS examines the angle between neural responses in feature space, and obtains an average angle for the data points closest to it. Intuitively, a small average angle among neighboring points implies that the data is tightly packed together into clusters, whereas larger angles suggest broadly distributed, unclustered data. We found that our neural responses were more tightly packed together than would be observed for an unclustered, Gaussian reference distribution (Suppl. Fig. S4).

Finally, we also compared the estimated the number of clusters in the data to those obtained using other clustering metrics, in particular the silhouette score and adjusted Rand Index (Suppl. Fig. S5). The silhouette score quantifies cluster quality with two components: it provides high scores when points within a cluster are close in feature space, and penalizes small inter-cluster distances that arise when clusters are nearby in feature space. The adjusted Rand Index (ARI) does not assess cluster quality through distances among datapoints in feature space, but instead is a measure of reproducibility of a clustering result. Specifically, it quantifies the probability that two labels from separate clustering results would arise by chance. We generated labels for ARI by subsampling our data (90% sampling without replacement) and calculated ARI on the subsampled dataset and the full dataset. ARI results demonstrated a broad range of possible numbers of clusters with reproducible cluster labels (Suppl. Fig. S5 A). In contrast, the silhouette score gave inconclusive results on our lOFC data (Suppl. Fig. S5 B). The dimensionality and covariance of our lOFC responses lies in a regime in which clusters are tightly packed in feature space, a regime in which silhouette score is known to underestimate the number of clusters in other contexts [11, 12]. To assess this more directly, we compared the results from different clustering metrics on artificial data with known statistics (deriving from 5 clusters). We found that in the data-like regime, the silhouette score substantially underestimated the optimal number of clusters (2 clusters, Suppl. Fig. S6, see Discussion and Methods). The gap statistic underestimated cluster number on synthetic data, but to a lesser degree. Taken together, these analyses suggest that the gap statistic is a conservative but principled metric for determining the number of clusters in the statistical regime of the lOFC data.

### 3.2 Neural responses in lOFC are well-captured by a generalized linear model that includes attributes of offered rewards, choice, and reward history

We next sought to characterize the encoding properties of neurons in each cluster. To that end, we modeled each neuron’s trial-by-trial spiking activity with a generalized linear model with Poisson noise (Fig. 3). This approach has been previously used in sensory areas to model the effects of external stimuli upon neural firing [13, 14], and more recently has been extended to higher-order areas of cortex during decision-making tasks [15]. In this model, the time-dependent firing rate (λ*_t_*) of a neuron is the exponentiated sum of a set of linear terms

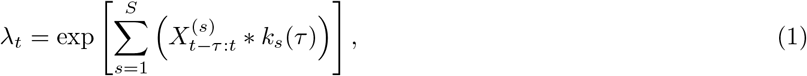

where the variables 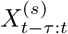 are the stimuli and behavioral events against which neural activity is regressed; *k_s_*(*τ*) denote the response kernels that are convolved with the task variables to model time-dependent responses to each variable (Fig. 3C). The probability that a given number of spikes would occur in a discrete time bin of Δ*t* = 50ms is given by a homogeneous Poisson process with mean λ*_t_*Δ*t*.

**Figure 3:**
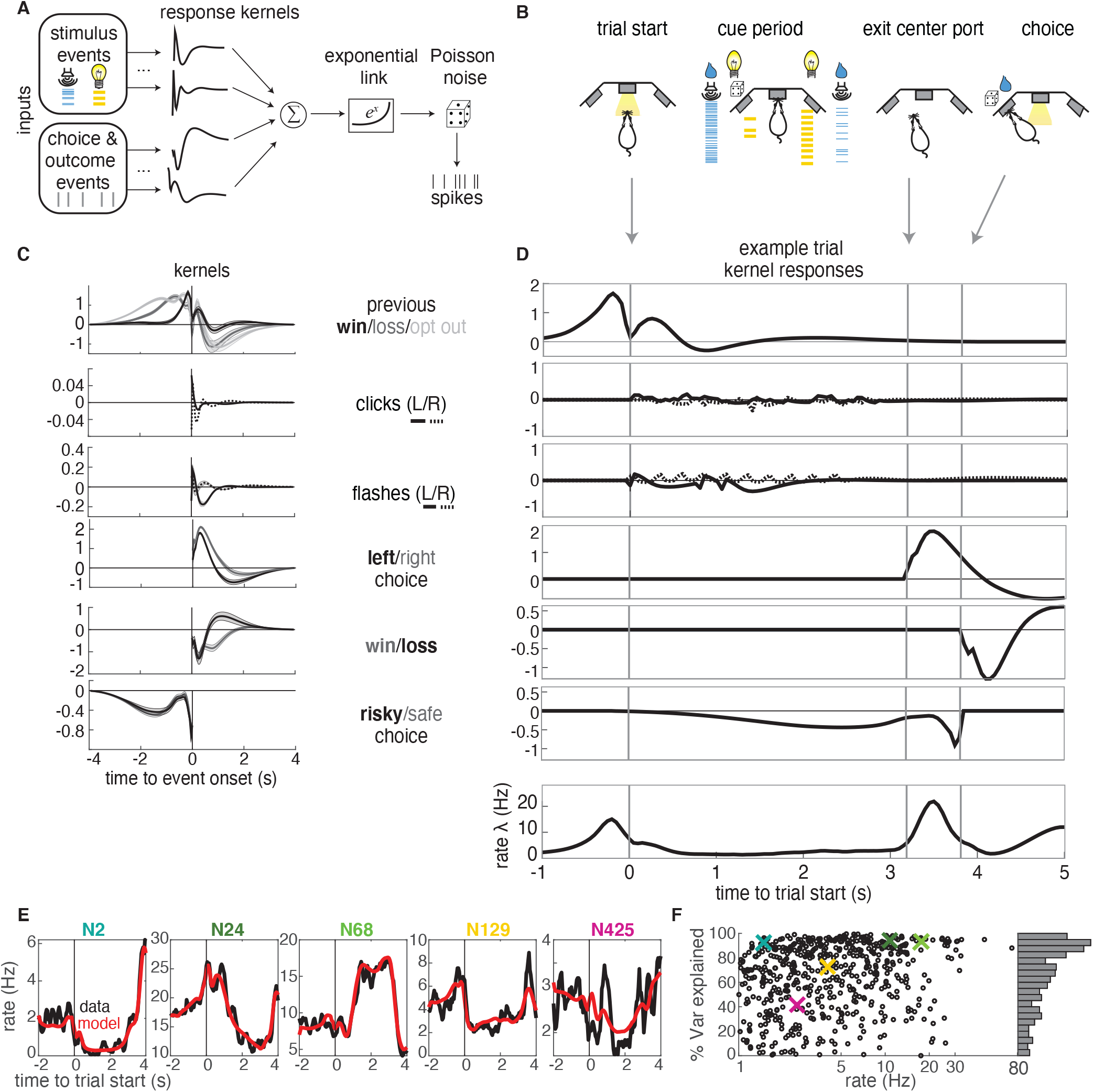
GLM analysis. A. Model schematic: the timings of external stimuli conveying reward attributes, as well as timings of choices and outcomes on each trial serve as model inputs. Nonlinear response kernels convolved with each input generates time dependent responses that are summed and exponentiated to give a mean firing rate, λ*_t_*, in each time bin. Spikes are generated from a Poisson process with mean firing rate λ*_t_*. B. Task schematic illustrating the key choice and outcome inputs to the model. C. Kernel fits for a sample neuron. Kernels are grouped by the aspects of the task that they model. Error bars denote estimated kernel standard deviation (Methods). D. Timing of each kernel’s contribution in an example trial. The kernels in bold from panel C are the kernels that are active in this trial. The resulting model-predicted firing rate is shown in the bottom row. E. Representative PSTHs to held-out testing data from 5 different neurons (black) and model prediction (red). F. Variance explained for each neuron, with sample neurons from E denoted by correspondingly colored crosses. The distribution of *R*^2^ values is presented along edge of the panel.

The task variables and example kernels for our model are shown for a sample neuron in Fig. 3C. Variables were binarized such that 1 (0) denoted the occurrence (absence) of an event. The task variables included reward (win or loss on the current or previous trial), choice (left or right), and the timing of cues indicating reward attributes for the left and right offers (reward volume conveyed by auditory clicks and reward probability conveyed by visual flashes). We also included the average reward rate prior to starting each trial and the location of the trial within the session (not shown). Given the large number of model parameters, we used a smooth, log-raised cosine temporal basis and L2 regularization to prevent overfitting (Methods). Based on model comparison, we found that including a spike history term as in other GLM approaches did not improve our model, presumably due to the fact that we are modeling longer timescale responses.

The chosen model parameters were selected by model comparison against several alternative models using crossvalidation. Model comparison favored a simpler binary win/loss representation of rewarded outcomes over richer representations of reward volume on the current or previous trial (Suppl. Fig. S7). The model reproduced the trial-averaged PSTHs of individual neurons (Fig. 3E). To quantify model performance, we calculated the proportion of variance explained (*R*^2^) for held-out testing data (Fig. 3F). The model captured a high percentage of variance for most of the neurons in our dataset. A small fraction of neurons exhibited negative *R*^2^ values (69 units, Suppl. Fig. S8), indicating that the model produced a worse fit of the test data than the data average. Our liberal inclusion criteria did not require neurons to exhibit task-modulation of their firing rates, so these neurons were likely not task-modulated, and were excluded from subsequent analyses.

### 3.3 Clustering reveals distributed encoding of most task variables across subpopulations of lOFC neurons

We next sought to characterize the extent to which neurons that belonged to different clusters might be “functionally distinct,” and encode different task-related variables [7, 8]. To address this question, we computed two complementary but independent metrics, both based on the GLM fit: the coefficient of partial determination and mutual information. The coefficient of partial determination (CPD) quantifies the contribution of a single covariate (e.g., left/right choice) to a neuron’s response by comparing the goodness-of-fit of the encoding model with and without that covariate. In other words, the CPD quantifies the amount of variance that is explained by kernels representing different aspects of the model. We computed the CPD in a sliding time window throughout the trial, and compared CPD values to a shuffle test, in which trial indices were randomly shuffled relative to the design matrix before generating model predictions. CPD values that were within the 95% confidence intervals of the shuffled distribution were set to 0 before averaging CPD values over neurons in a cluster. The average CPD values for different task variables is shown for clusters based on each clustering procedure in Fig. 4A and Fig. 4C. Note that CPD plots are aligned to different events, depending on the covariate.

**Figure 4:**
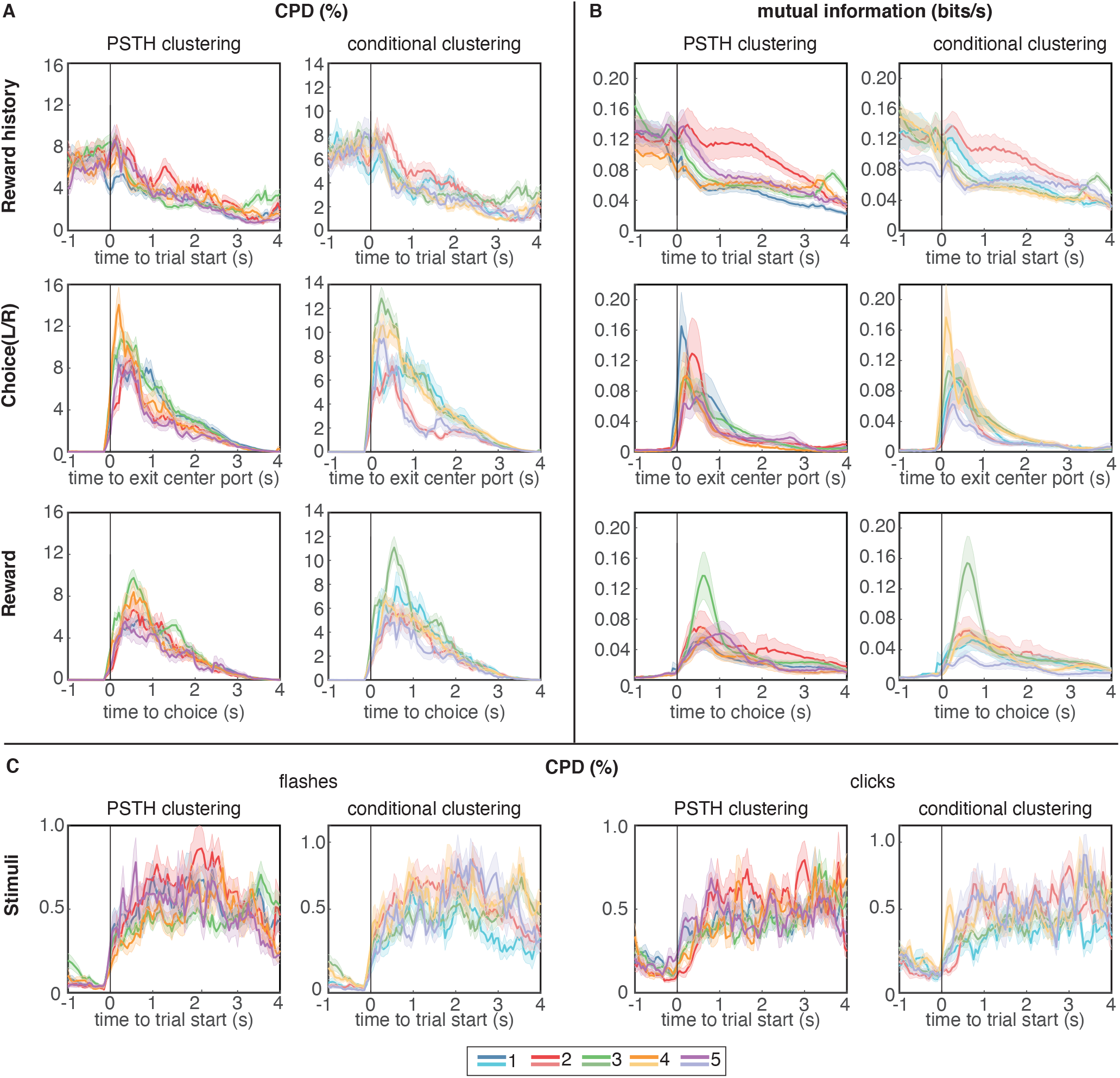
Coefficient of partial determination (CPD) and mutual information (MI) reveal broad encoding of all task parameters in each cluster. A. CPD for choice and reward outcome encoding: Reward history panels quantify encoding of outcome on the previous trial (previous win/loss/opt out), choice panels convey encoding of choices on the current trial (left/right), and reward panels quantify encoding of outcomes on the current trial (win/loss). B. Mutual information for same task parameters. C. CPD of flashes (left panels) and clicks (right panels) that encode reward probability and reward volume, respectively. CPD and MI values are averaged over neurons in each cluster; error bars show s.e.m. Results for PSTH clustering are in the left columns, and results for conditional clustering are in the right columns.

We also computed the mutual information (MI) between neural spike trains and different model covariates (Methods). Our approach relates the statistics of trial-level events to firing rates, which allows us to assess the information content for each stimulus throughout the entire time course of a trial. Such an approach does not easily generalize to the statistics of within-trial events such as the information represented in stimulus clicks and flashes, so we restricted the MI analysis to the other covariates: reward history, choice, and reward outcome. The average MI for these variables is shown for clusters based on the trial-averaged PSTHs in Fig. 4B.

In general, regardless of the metric (CPD or MI) or clustering procedure (PSTH clustering or conditional clustering), task variables appear to be broadly encoded across neural subpopulations, with similar temporal dynamics and average CPD/MI values across clusters. All clusters encoded cues conveying reward volume and probability during the cue period (Fig. 4C), although it is worth noting that the magnitude of CPD for clicks and flashes was an order of magnitude lower than for the other task variables. Therefore, encoding of cues representing reward attributes was substantially weaker than encoding of reward history, choice, and outcome. It may be surprising that neurons with strikingly different PSTHs appear to encode task variables with similar time courses. However, PSTHs marginalize out all conditional information, so a PSTH carries no information about encoding of task or behavioral variables, *per se*. Additionally, to assess whether different clusters may have a different composition across neuron types, we classified single units based on their waveforms into regular spiking units with broad action potential (AP) and after-hyperpolarization (AFP) widths, or fast-spiking units with narrow AP and AHP widths. We found the distribution of these two unit types to be similar across clusters (Suppl. Fig. S10).

Reward history was most strongly encoded at the time of trial initiation and decayed over the course of the trial, consistent with previous analyses [6]. Neurons belonging to cluster 2, which strongly overlap in both clustering procedures (Fig. 2E), seem to exhibit slightly more pronounced encoding of reward history during the trial, compared to the other clusters, although these neurons were also persistently active during this time period. Encoding of choice and reward outcomes were phasic, peaking as the animal made his choice and received outcome feedback, respectively, and were broadly distributed across clusters (Fig. 4A-B).

### 3.4 Reward history re-emerges before choice in a distinct subset of lOFC neurons

The CPD and MI analysis revealed one aspect of encoding that is unique to cluster 3. Both clustering methods identified largely overlapping populations of neurons in this cluster (Fig. 2E), indicating that these neurons exhibited similar temporal response profiles as well as encoding. Like neurons in the other clusters, neurons in cluster 3 encoded reward history at trial initiation, and that encoding decreased through the cue period. However, for neurons in this cluster, encoding of reward history re-emerged at the time preceding the animal’s choice on the current trial (Fig. 5A-B, green lines). This “bump” of reward history encoding late in the trial was unique to cluster 3 regardless of the clustering method, and was observable in both CPD and MI (Fig. 4A-B). This result was further corroborated by computing the average discriminability index or (*d*′) for reward history, which is a model-agnostic metric that quantifies the difference in mean firing rate in each condition, accounting for response variance over trials. Cluster 3 was unique from the other clusters for having a subset of neurons with high *d*′ values for reward history at the time preceding the animal’s choice (Fig. 5C). We additionally found that encoding of previous wins and losses during this time period only extended to the previous trial, and did not encode the outcome of additional past trials (Suppl. Fig. S11). Clustering of PSTHs aligned to later events in the trial showed more detailed structure of this late-in-trial encoding of reward history, with some neurons encoding the previous trial just before the choice, and others at the time of choice (Suppl. Fig. S12 C). We note that clustering on PSTHs aligned to later events produced similar results, in terms of the number of clusters and the within-cluster responses (Suppl. Fig. S12 B)

**Figure 5:**
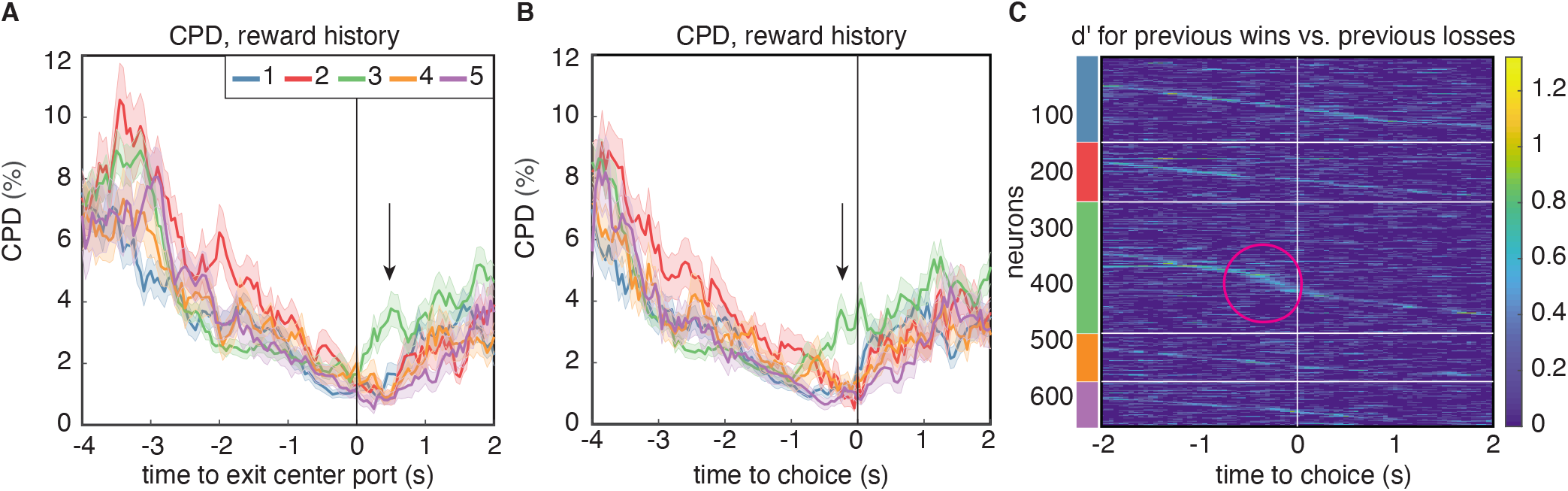
Encoding of reward history emerges late in the trial for cluster 3. A. CPD for reward history, aligned to leaving the center poke after the cue period. Note the peak in encoding that is isolated to cluster 3 (black arrow). B. CPD similarly aligned to choice, demonstrates that this encoding occurs before reward delivery on the current trial. C. Sensitivity, *d*′, across time and neurons for reward history has activity isolated to cluster 3 (magenta circle). *d*′ is sorted within the PSTH-based clusters by time-to-peak.

Notably, cluster 3 also exhibited the most prominent encoding of reward outcome compared to the other clusters (Fig. 4A-B, green, bottom row). This suggests that this subset of neurons may be specialized for representing or even integrating information about reward outcomes on previous and current trials. We wondered if this might reflect adaptive value coding, in which value representations in OFC have been observed to dynamically reflect the statistics of offered rewards [16–21]. Adaptive value coding, which is thought to reflect a divisive normalization or range adaptation mechanism, allows the circuit to efficiently represent values in diverse contexts in which their magnitudes may differ substantially. As such, it provides key computational advantages, such as efficient coding, or the maximization of mutual information between neural signaling and the statistics of the environment [17–20, 22–25].

According to divisive normalization models of subjective value, the value of an option or outcome is divided by a recency-weighted average of previous rewards [17, 24, 25]. Therefore, if neurons in OFC exhibit adaptive value coding, we would predict that they would exhibit stronger reward responses following unrewarded trials, and weaker responses following rewarded trials (Fig. 6C). Put another way, regressing the firing rate against current and previous rewards should reveal coefficients with opposite signs [26]. Neurons with regression coefficients for current and previous rewards with the same sign would be modulated by current and previous rewards, but not in a way that is consistent with the adaptive value coding hypothesis (Fig. 6D).

**Figure 6:**
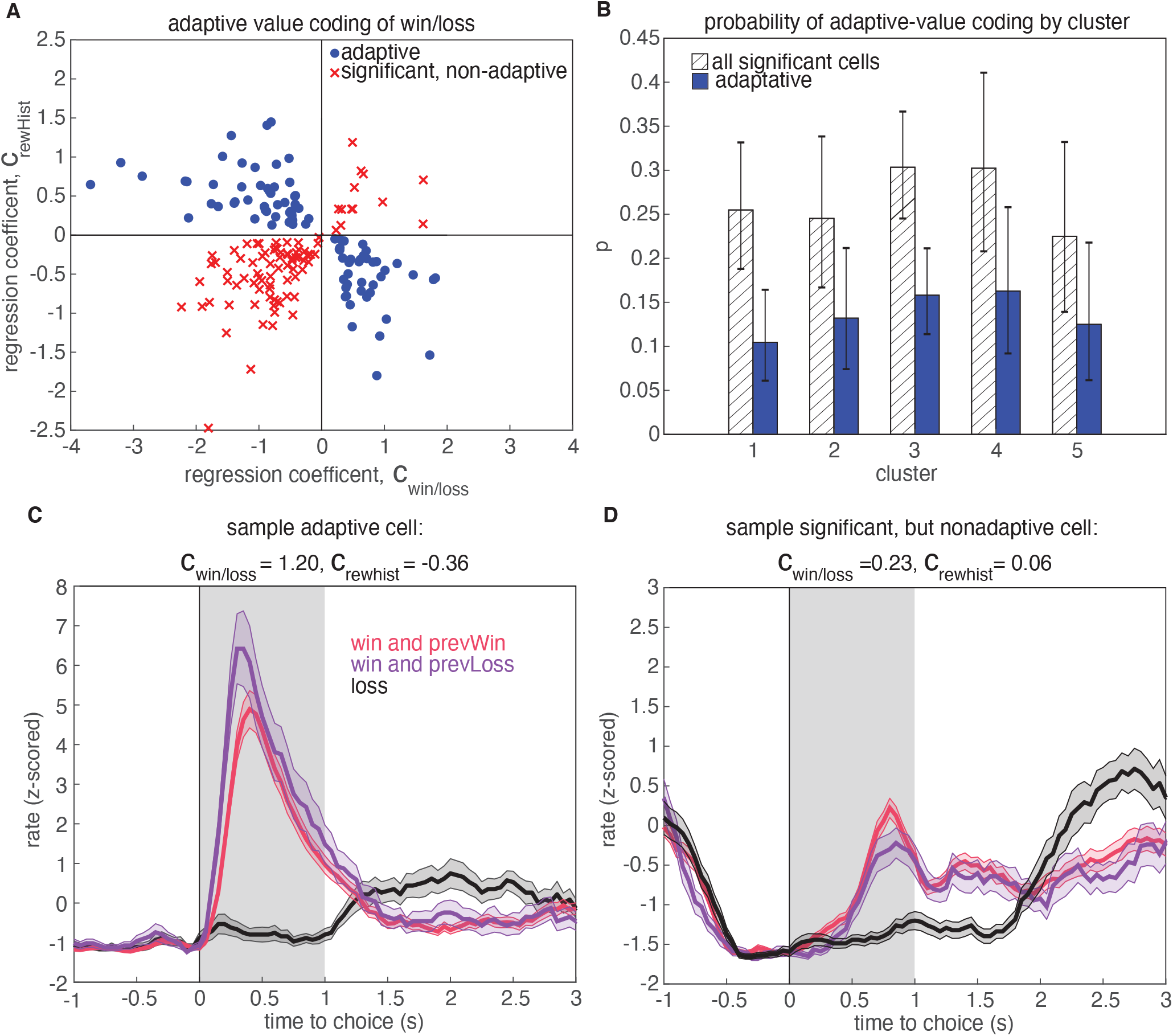
Adaptive value coding analysis using linear regression of firing rates for the 1s period following reward delivery against current and previous reward outcomes. A. Regression coefficients for the current reward outcome, *c*_win/loss_ (parameterized as win=1, and loss=-1), and previous trial outcome, *c*_rewhist_. 180 neurons had significant coefficients for both regressors (*p* < 0.05, t-test), and 81 neurons had coefficients with opposite signs, consistent with adaptive value coding (blue dots). The remaining neurons have differential responses due to reward history, but inconsistent with adaptive value coding (red crosses). B. Probability that a model with significant regressors for both current and past reward outcome would come from a given cluster. Shaded regions denote all models from panel A, and blue bars show the probability for adaptive neurons only. Error bars are the 95% confidence interval of the mean for a binomial distribution with observed counts from each cluster. C. Example cell demonstrating adaptive value coding. Shaded gray region denotes time window used to compute mean firing rate for the regression. D. Sample cell demonstrating significant modulation due to reward history, but with a relationship inconsistent with adaptive value coding.

To test this hypothesis, we regressed the firing rates of neurons in a 1 second window after reward receipt against reward win/loss outcomes on the current trial, as well as the win/loss outcome from the previous trial (Fig. 6, Methods). We found that 180 neurons (27% of population) exhibited significant coefficients for both current reward and reward history regressors, and 45% of these units demonstrated adaptive value coding indicated by the opposite signs of regression coefficients for current and previous reward outcomes (81 units, Fig. 6A, blue). To determine if these adaptive neurons preferentially resided in a particular cluster, we calculated the cluster-specific probability of a neuron demonstrating either significant coefficients for rewarded volume and reward history, or having coefficients with opposite signs, consistent with adaptive value coding (Fig. 6B). We found that no cluster preferentially contained adaptive units with higher probability (comparison of 95% binomial confidence intervals). We also investigated whether adaptive value coding was present for rewarded volume representations, and similarly found that neurons did not preferentially reside in any one cluster (30 adaptive units out of 79 significant units, Suppl. Fig. S14). Notably, activity during the reward epoch did not appear to encode a reward prediction error (Suppl. Fig. S13), defined as the difference between the expected value and the outcome of the chosen option, as has been previously reported in rat and primate OFC [26–30].

### 3.5 Neurons in cluster 3 have similar responses to previously reported striatum projection neurons

Studies of OFC indicate that long-range projections to different downstream circuits exhibit distinct encoding and/or make distinct contributions to behavior [7, 8, 31]. For instance, previous work from [7] used optogenetic tagging to record from OFC neurons that project to the ventral striatum, and found that these cells exhibited responses that seemed to correspond to a distinct cluster with stereotyped responses. This cluster encoded the trial outcome just after reward delivery, with larger responses following unrewarded trials, and also encoded the negative integrated value of the chosen option during the inter-trial interval, until the start of the next trial (Fig. 7A). We examined the average PSTHs of each cluster aligned to trial outcome and the start of the next trial, to see if any cluster resembled the corticostriatal responses described in [7]. Specifically, we plotted the cluster-averaged firing rates for trials classified as a loss, a low-reward trial (6 or 12 *μ*l), or a high-reward trial (24 or 48 *μ*l) (Fig. 7 B). We found that several clusters prominently encoded the reward outcome after reward (clusters 1, 3, 5); however, only cluster 3 encoded the negative (i.e., inverted) value of the reward during the inter-trial interval until the start of the next trial. Therefore, one of the clusters that we identified in our data exhibits qualitatively similar responses to the corticostriatal projection neurons identified by clustering in [7].

**Figure 7:**
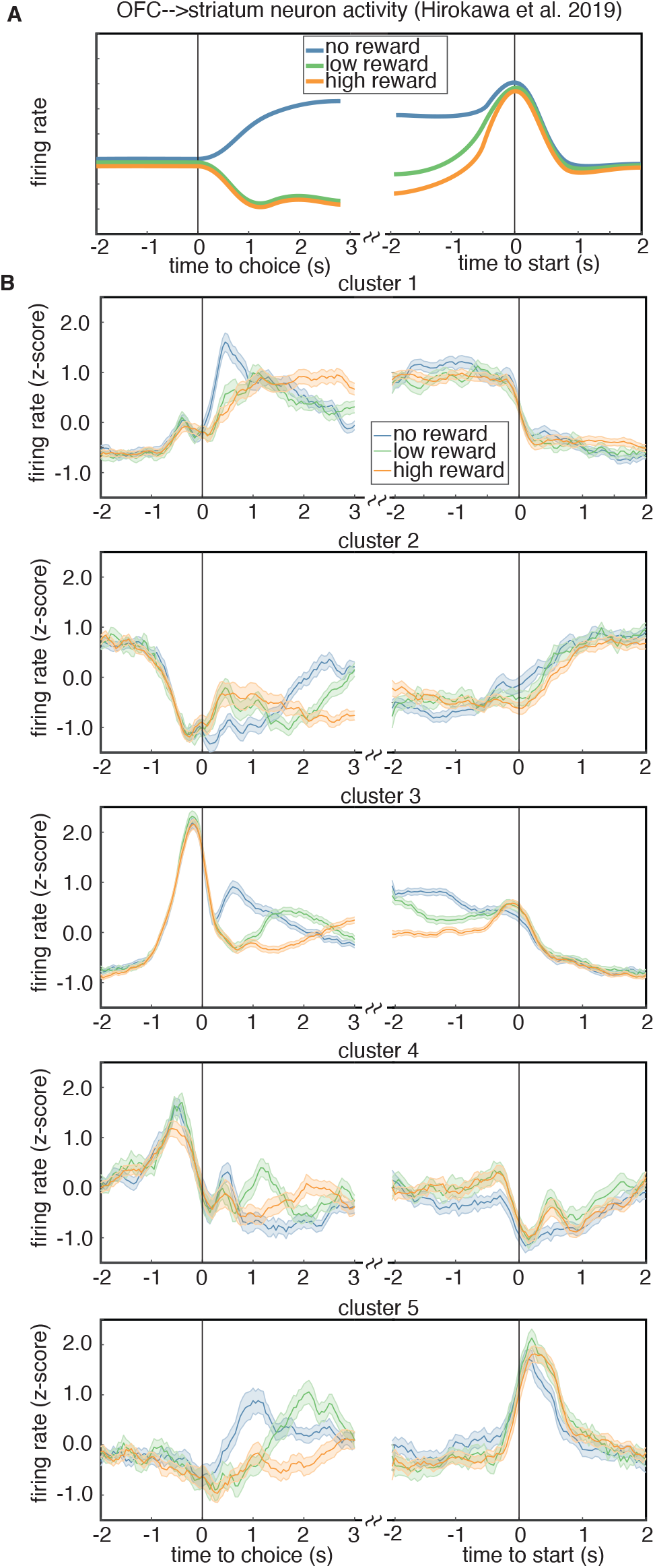
A. Schematic of the encoding properties of a cluster of putative corticostriatal projection neurons from [7]. Neurons from this cluster encode the outcome of a trial just after choice, as well as negative (*i.e*., inverted) integrated value during the intertrial interval, until the start of the next trial. B. Cluster-averaged response to rewarded volume magnitude. Each sub-panel shows the average response to either a trial loss (blue), a low reward (6 or 12 *μ*l, green), or a high reward (24 or 48 *μ*l, orange).

## 4 Discussion

We analyzed neural responses in lOFC from rats performing a value-based decision-making task, in which they chose between left and right offers with explicitly cued reward probabilities and volumes. Despite the apparent response heterogeneity in our dataset, two independent clustering methods revealed that neurons belonged to one of a small number of distinct clusters. We clustered based on the marginalized PSTHs over all trials, and found that subpopulations of neurons exhibited characteristic temporal response profiles. To our knowledge, using temporal responses that do not explicitly contain conditional information is a novel approach for identifying distinct neural response profiles in prefrontal cortex. We also clustered based on a conditional feature space for each neuron, consistent with previous work [7, 8]. The feature space for each neuron corresponded to its average firing rate on different trial types, in select time windows that most closely corresponded to the differentiating covariate on each trial (e.g., wins vs. losses at the time of reward feedback). Notably, these independent clustering methods identified highly robustly identifiable groups of neurons for two of the clusters, indicating that neurons in these clusters can be identified by either thei temporal response profiles or their task variable encoding, as defined by the feature space.

Previous studies that clustered using a conditional feature space identified a larger number of clusters than we did here [7, 8]. This could reflect the different metrics used to select the number of clusters (the gap statistic in this study; compared to PCA and silhouette scores [8], or adjusted Rand index [7]). Another difference is the time window that was used to generate the conditional feature space. In [7], for instance, the authors used the same time window for each feature, which was after the rat made a choice but before it received feedback about the outcome of that choice. There was no such epoch in our behavioral paradigm, precluding a more direct comparison. However, a principled choice of clusters using the gap statistic still provided a useful tool for investigating the encoding of task variables in lOFC.

### 4.1 Rat lOFC weakly encodes reward attributes

Our task design allowed us to isolate neural responses to sensory cues that conveyed information about distinct reward attributes, because these cues were presented independently and variably in time. However, our GLM revealed weak encoding of these reward attributes – reward probability and volume – across all clusters, regardless of the clustering method. This was observable by examination of the relative magnitude of the flash and click kernels in individual neurons, and also by the CPD metric. The average flash and click CPD values were an order of magnitude smaller than for the other covariates, indicating that flashes and clicks did not contribute substantially to neural firing. Therefore, while behavioral analyses have shown that rats used the flashes and clicks to guide their choices [5], these cues were not strongly represented in lOFC. This weak encoding of reward attributes, whose combination would specify the subjective value of each offer [5], is potentially consistent with a recent study of rat lOFC during a multi-step decisionmaking task that enabled dissociation of choice and outcome values [30, 32]. Recordings in that study revealed weak encoding of choice values, but strong encoding of outcome values, and optogenetic perturbations suggested a critical role for lOFC in guiding learning but not choice [30]. Other studies in rat lOFC have reported strong encoding of reward and outcome values specifically following action selection, but not preceding choice [7, 30, 33–36]. However, recordings from rat lOFC in sensory preconditioning paradigms have revealed cue-evoked responses that may reflect inferred value [29, 37]. Studies in non-human primates have also reported strong encoding of offer values in OFC after presentation of a stimulus, and before choice [26, 38–40]. A recent study in mouse OFC used olfactory cues to convey reward attributes (juice identity and volume) and reported encoding of offer values before choice [41]. It is unclear what factors may account for the pronounced encoding of offer value before choice in [41], but minimal encoding of offer value before choice in our data. One important difference is that the cues that conveyed reward attributes in the present study needed to be integrated over time. The integration process itself likely does not occur in OFC [39]. It is possible that OFC would only represent the offer value once the animal had accumulated enough sensory evidence to infer it. If that inference occurred at a different time on each trial, depending on the strength of the sensory evidence and stochasticity of the accumulation process, then representations of offer value would not be observable in the trial-averaged neural response.

### 4.2 Adaptive coding in rat lOFC

Previous studies in primate medial [18] and central-lateral [19, 20, 26, 42] OFC have reported subsets of neurons that adjust the gain of their firing rates to reflect the range of offered rewards, a phenomenon referred to as adaptive value coding. This type of coding is efficient because it would allow OFC to accurately encode values in diverse contexts that may vary substantially in reward statistics [17], analogous to divisive normalization in sensory systems [22, 23].

According to divisive normalization models of subjective value, the value of an option is divided by a recency-weighted average of previous rewards [17, 24, 25]. Therefore, we would predict that neurons implementing adaptive value coding would exhibit stronger responses following unrewarded trials, and weaker responses following rewarded trials. Consistent with this hypothesis, we identified a subset of neurons that had significant coefficients for rewards on current and previous trials, with opposite signs, and while this fraction was somewhat modest (~15%), it was comparable to the proportion of adaptive value coding neurons observed in central-lateral primate OFC [26]. Notably, we did not find any evidence of encoding of a reward prediction error during this epoch, or the difference between the expected value of the chosen lottery and the outcome. Therefore, the differential encoding of reward outcomes depending on reward history reflected a discrepancy between reward outcomes and recent experience, not a discrepancy between reward outcomes and expectations (as in a reward prediction error). These were dissociable in our task, due to the sensory cues that explicitly indicated expected value on each trial. We emphasize that adaptive value coding is likely not a specialization of lOFC, but probably occurs broadly in brain areas that represent subjective value.

### 4.3 Mixed selectivity in OFC

The complexity and diversity of responses found in the prefrontal cortex, including the OFC, has led to questions about whether or not neurons with mixed selectivity or specialized responses represent behavioral and cognitive variables [7, 8, 10, 43, 44]. We tested if OFC neural responses belonged to clusters in our task using two separate methods: the gap statistic, and the PAIRS statistic. Both methods confirm that there is statistically significant clustering in our data. We emphasize, however, that mixed selectivity is fundamentally a property of individual neurons. We did not directly investigate whether individual lOFC neurons in our dataset exhibited mixed selectivity, but instead focused on encoding at the level of clusters. The broad encoding of task variables in each cluster suggests that lOFC neurons probably exhibit mixed selectivity, but a formal assessment was beyond the scope of our study.

### 4.4 Representations of reward history in OFC

In studies that require animals to learn and update the value of actions and outcomes from experience (*i.e*., in accordance with reinforcement learning), values are often manipulated or changed over the course of the experiment to assess behavioral flexibility, value-updating, and goal-directed behavior. In rodents and primates, lesion and perturbation studies have shown that the OFC is critical for inferring value in these contexts, suggesting an important role in learning and dynamically updating value estimates for model-based reinforcement learning [30, 32, 45–50]. Neural recordings in these dynamic paradigms have revealed activity in OFC that reflects reward history and outcome values, which could subserve evaluative processing and learning [30, 33, 51].

Other studies, including this one, have used sensory cues (*e.g*., static images, odors) to convey information about reward attributes. Once learned, the mapping between these cues and reward attributes is fixed, and the subject must choose between options with explicitly cued values. Neurons in the central-lateral primate OFC and rat lOFC have been shown to represent the values associated with these sensory cues in their firing rates, as well as reward outcomes and outcome values [7, 33, 37–39, 42, 52–57]. However, the extent to which OFC is causally required for these tasks is a point of contention, and may differ across species [41, 58, 59]. Notably, even when reward contingencies are fixed over trials, animals often show sequential learning effects [5, 60], and reward history representations in OFC have been reported in tasks with stable reward contingencies [6, 26, 61].

In this study, we have described dynamic trial-by-trial changes in firing rates that reflected reward history just preceding the choice. These reward history representations might influence ongoing neural dynamics supporting the upcoming choice [62], and/or mediate learning. In contrast to broadly distributed adaptive value coding, this activity was restricted to a particular subset of neurons that were identifiable by two independent clustering methods, and that exhibited the strongest encoding of reward outcomes. Intriguingly, the responses of neurons in this cluster (cluster 3) are qualitatively similar to the responses of identified corticostriatal cells in [7], raising the possibility that cluster 3 neurons also project to the striatum. lOFC neurons that project to the striatum likely mediate learning: ablating OFC projections to the ventral striatum during a reversal learning task demonstrated that this projection specifically supports learning from negative outcomes [31].

In reinforcement learning accounts of basal ganglia function, it is thought that cortical inputs to the striatum encode information about the state the animal is in, and corticostriatal synapses store the learned values of performing actions in particular states in their synaptic weights (*i.e*., Q-values), which can be updated and modulated by dopamine-dependent plasticity [63, 64]. Coincident activation of cortical inputs and striatal spiking can tag corticostriatal synapses for plasticity in the presence of dopamine [65]. One reason that lOFC neurons might encode reward history at the time of choice is so that contextual inputs reflecting the animal’s state would include its recent reward history. We have previously shown that optogenetic perturbations of lOFC in this task, triggered when rats exited the center port, disrupted sequential trial-by-trial learning effects [6]. These sequential learning effects are not optimal for decision making in this task, as trials were independent and contained explicitly cued reward contingencies. Suboptimal sequential biases are observed across species and domains [66–68], and reflect an innate difficulty in judging true randomness in an environment [69]. The neural circuits supporting these sequential biases could potentially derive from the subpopulation of lOFC neurons in cluster 3, which may mediate the effect of previous trials on the animal’s behavior either by influencing lOFC dynamics supporting the upcoming choice [62], or via projections to the striatum that mediate learning.

## 5 Methods

### 5.1 Animal subjects and behavior

The details for the animal subjects, behavioral task, and electrophysiological recordings have been described in detail elsewhere [5, 6]. Briefly, neural recordings from three male Long-Evans rats were used in this work. Animal use procedures were approved by the Princeton University Institutional Animal Care and Use Committee and carried out in accordance with National Institutes of Health standards.

Rats were trained in a high-throughput facility using a computerized training protocol. The task was performed in operant training boxes with three nose ports, each containing an LED. When the LED from the center port was illuminated, the animal was free to initiate a trial by poking his nose in that port (trial start epoch). While in the center port, rats were continuously presented with a train of randomly timed auditory clicks from a left and right speakers. The click trains were generated by Poisson processes with different underlying rates [70]; the rates from each speaker conveyed the water volume baited at each side port. Following a variable pre-flash interval ranging from 0 to 350 ms, rats were also presented with light flashes from the left and right side ports, where the number of flashes conveyed reward probability at each port. Each flash was 20 ms in duration, presented in fixed bins, and spaced every 250 ms to avoid perceptual fusion of consecutive flashes. After a variable post-flash delay period from 0 to 500 ms, there was an auditory “go” cue and the center LED turned back on. The animal was then free to leave the center port (exit center port epoch) and choose the left or right port to potentially collect reward (choice epoch).

### 5.2 Electrophysiology and data pre-processing for spike train analyses and model inputs

Tetrodes were constructed from twisted wires that were either PtIr (18 μm, California Fine Wire) or NiCr (25 μm, Sandvik). Tetrode tips were platinum- or gold-plated to reduce impedances to 100–250 kΩ at 1 kHz using a nanoZ (White Matter LLC). Microdrive assemblies were custom-made as described previously [71]. Each drive contained eight independently movable tetrodes, plus an immobile PtIR reference electrode. Each animal was implanted over the right OFC. On the day of implantation, electrodes were lowered to 4.1 mm DV. Animals were allowed to recover for 2–3 weeks before recording. Shuttles were lowered 30–60 μm approximately every 2–4 days.

Data were acquired using a Neuralynx data acquisition system. Spikes were manually sorted using MClust software. Units with fewer than 1% inter-spike intervals less than 2 ms were deemed single units. For clustering and model fitting, we restricted our analysis to single units that had a mean firing rate greater that 1 Hz (659 units). To convert spikes to firing rates, spike counts were binned in 50 ms bins and smoothed using Matlab’s smooth.m function with a 250ms moving window. Similarly, our neural response model fit spike counts in discretized bins of 50ms. When parsing data into cross-validated sets of trials balanced across conditions, a single trial was considered as the window of [−2,6]s around trial start. In all other analyses, conditional responses were calculated on trials with a window [−2,4]s for data aligned to trial start, and [−4,4]s for choice-aligned or leave centerpoke-aligned responses.

### 5.3 Clustering of lOFC responses

#### 5.3.1 Feature space parametrization

To analyze the heterogeneity in time-dependent lOFC responses, our first clustering procedure utilized trial-averaged PSTHs from each neuron to construct the feature space for clustering. Specifically, for a set of *N* neurons and trials of *T* timepoints, we z-scored the PSTH of each neuron and combined all responses into a matrix 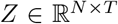, then performed PCA to obtain principal components, *W*, and score, *M*, as *Z* = *MW^T^*. We found that *k* =18 components explained > 95% of the covariance in *Z* and used the first *k* columns of *M*, the PSTH projected onto the top *k* PC, as our feature space on which to perform clustering.

Our second clustering procedure used time-averaged, conditional neural responses to construct the feature space for k-means clustering. This is similar in form to the approach in [7]. The feature space consisted of its z-scored firing rate conditioned on choice, reward outcome, reward history, presented offer value (EV of left and right offers), and rewarded volume, in time bins that most often corresponded to differential encoding of each variable (Fig. 2A, Fig. 2B). Specifically, we used conditional PSTHs that depend on a single condition, and marginalized away all other conditions. The conditions, *X*^(*M*)^ = *x_j_*, are grouped into three categories for our task. Choice and outcome information is **X**^(reward)^ ∈ {win,loss}, **X**(^rewardHistory^) ∈ {previous Win,previous loss}, **X**^(choice)^ ∈ {left,right}. Presented offer attributes on left and right ports are the expected value of reward as *EV* = *pV*, where *p* is reward probability conveyed through flashes, and *V* is the volume offer conveyed through Poisson clicks. Values were binned on a log-2 scale: **X^(EVL)^**, **X^(EVR)^** ∈ {[0, 6), [6, 12), [12, 24), [24, 48]*μ*l}. Rewarded value is **X**^(rewVol)^ ∈ {0, 6, 12, 24, 48 *μ*l}.

The conditional PSTH responses were z-scored, and then each conditional PSTH was averaged over the time window in which the behavioral variable was maximally encoded (dictated by peak location in CPD, see Methods below) to obtain a conditional firing rate as a single feature for clustering. Specifically, the time windows for reward and reward volume information were averaged over [0, 3]s after reward delivery, [—1, 2]s after trial start for reward history, [0, 1.5]s after exiting the center port for left/right choice, and [–1, 0]s before the animal’s choice (*i.e*., entering the side port) for expected value of presented offers. These 19 features were combined and pre-processed using PCA in the same way as the PSTH-based clustering to yield 11 features.

#### 5.3.2 Evaluation of k-means cluster quality: gap statistic

For each clustering procedure, we utilized k-means clustering to locate groups of functionally distinct responses in lOFC, and used the gap statistic criterion to determine a principled choice of the best number of clusters (evalclusters.m in Matlab) [9]. Specifically, we locate the largest cluster size *K* for which there was a significant jump in gap score *Gap*(*K*),

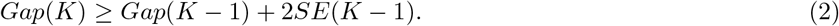

This is similar to a standard option in evalclusters.m (‘SearchMethod’= ‘firstMaxSE’), which finds the smallest instance in which a non-significant jump in cluster size is located. The two methods often agree. Finally, we used 5000 samples for the reference distribution to ensure convergence of results in the gap statistic.

#### 5.3.3 Alternative cluster evaluation metrics: adjusted Rand index (ARI) and silhouette score

The silhouette score calculates a weighted difference between inter- and intra-cluster distances. It yields high values for data that are tightly packed in the feature space and also far away from neighboring clusters. Intra-cluster distances for a data point *i* quantify the similarity of data within a cluster as

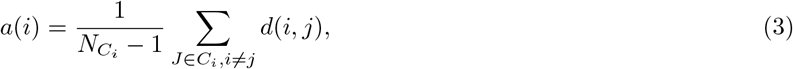

where *d*(*i,j*) is the Euclidean distance between points *i* and *j*, and *N_C_i__* is the number of data points in cluster *C_i_*. Similarly, the inter-cluster distance of data point *i* to data points of the closest neighboring cluster is given by

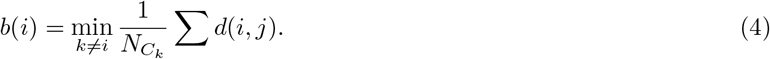

The silhouette score is given by

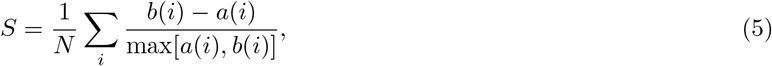

where *N* is the number of data points. Silhouette scores were calculated in python using sklearn.metrics.silhouette_score with the default options (*i.e*, metric=‘euclidean’).

The adjusted Rand index is a measure of reproducibility of a dataset. In this context, it calculates the probability that two cluster labelings on related data would arise by chance. We calculated ARI in python using sklearn.metrics.adjusted_rand_score. To estimate statistics, we calculated ARI between a set of labels generated from our full dataset and a sub-sampled dataset for each cluster number *K*. We sampled 90% of the population without replacement, and generated 100 such datasets to get a distribution of ARI values for each value of *K*.

To study the behavior of the silhouette score in the same data regime as the lOFC neural data, we generated ground-truth datasets of *N* = 500 data points with different ratios of within-cluster and total data covariance. In particular, we generated datasets with a K=5 clusters (100 data points per cluster) that had the same dimensionality (D=121) and the same total data covariance as the neural data. These clusters had within-cluster covariance that matched the average within-cluster covariance of the lOFC data. We then preprocessed this surrogate data in the same manner as our true data by projecting onto the PC components capturing > 95% variance (total-data covariance and average within-cluster covariance shown in Suppl. Fig. S6 A-B), and performed silhouette score analysis in this dimensionality-reduced subspace. We investigated different ratios of within-cluster and across-cluster by scaling the total-data covariance by a constant factor (0.1, 0.5, 1, 2, 3, 5) to determine to what extent the silhouette score can detect clusters that are tightly packed in the feature space. To gather statistics for each ratio of variances, we generated 100 datasets with random cluster means (drawn from a multivariate normal distribution) and plotted the mean ± s.e.m silhouette scores Suppl. Fig. S6 B). As a comparison, we also performed the gap statistic analysis on the same datasets (Suppl. Fig. S6 C). Finally, we visualized the groupings of these clusters by nonlinear projections (tSNE) in a two-dimensional feature space (Suppl. Fig. S6 D).

#### 5.3.4 Cluster labeling consistency and cluster similarity

To compare the consistency of results between the PSTH clustering and conditional clustering (Fig. 2E), we calculated *P*(*C*_conditional_|*C*_PSTH_), the conditional probability of a neuron being assigned to cluster *C*_conditional_ from the conditional clustering procedure, given that it was assigned to cluster *C*_PSTH_ in the PSTH cluster procedure. We evaluated the similarity of clusters within a given clustering procedure in two ways. First, we performed TSNE embedding of the features space to visualize cluster similarity in two dimensions (sklearn.manifold.TSNE in python, perplexity=50, n_iter=5000) [72]. We then colored each sample in this 2-D space based on cluster identity (Suppl. Fig. S3 A-B). We quantified the distance amongst clusters by calculating the cluster averaged Mahalanobis distance to the other clusters [73]. The Mahalanobis distance *D_M_*(*x*) calculates the distance of a sample *x_A_* to a distribution *B* with a known mean, *μ_B_*, and covariance, *S_B_*:

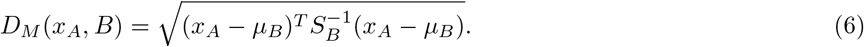

The cluster-averaged distances in Suppl. Fig. S3 C-D average *D_M_* over all samples from a given cluster *A* as

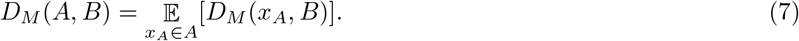

#### 5.3.5 PAIRS statistic

To test the null hypothesis that the data is not clustered at all, we followed the approach provided in [10]. We used our pre-processed PSTH and conditional feature spaces that were dimensionality reduced using PCA, and further processed them with a whitening transform so that the data had zero mean and unit covariance. For each data point, we estimated the average angle with *k* of its closest neighbors, *θ*_data_. We then compared the data to independent draws from a reference Gaussian distribution 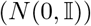, with the same number of data points and the same dimensionality as our data. To gather statistics, we generated *N* = 10,000 of these null model datasets, and aggregated the estimated angles 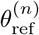 into a grand reference distribution. We chose our number of nearest neighbors *k* for averaging by locating the *k* such that our reference distribution had a median angle greater than π/4, following the approach in [10]. Distributions of angles between neighboring datapoints are presented in Suppl. Fig. S4. We further quantified the similarity of these two distributions through a Kolmogorov Smirnov test.

PAIRS is a summary statistic of the entire population, using the median from the data distribution, 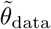, and the median of the grand reference distributions 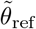,

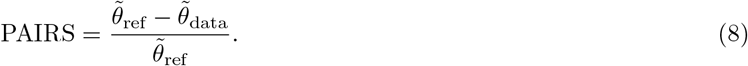

Reference PAIRS statistics were generated for each of the reference data sets, and statistical significance of the PAIRS statistic was assessed by calculating the two-sided p value for the data PAIRS compared to the distribution of reference PAIRS values. We assumed a normal distribution for the PAIRS reference values.

### 5.4 Generalized linear model of neural responses in lOFC

Our neural response model is a generalized linear model with an exponential link function that estimates the probability that spiking from a neuron will occur in time bin *t* with a rate λ*_t_*, and with Poisson noise, given a set of time-dependent task parameters. Specifically, for a design matrix **X** with *S* columns of task variables 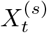, the probability of observing *y_t_* spikes from a neuron in time bin *t* + Δ*t* is modeled as

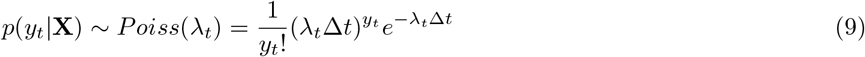

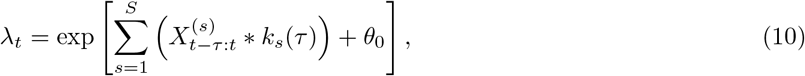

where λ*_t_* is the rate parameter of the Poisson process. Task variables are linearly convolved with a set of response kernels *k_s_*(*τ*) that allow for a time-dependent response from a neuron that may occur after (causal) or before (acausal) the event in the trial. The kernels are composed of a set of *N_s_* basis functions, 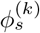, linearly weighted by model parameters 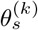:

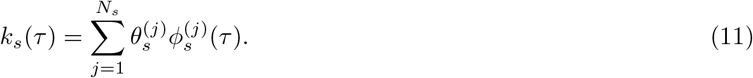

Additionally, we include a parameter *θ*_0_ that captures the background firing rate of each neuron.

We used 15 time-dependent task variables in our model that indicate both the timing of an event in the task, as well as behavioral or conditional information. Parameterization of the model in this manner with a one-hot encoding of each condition per variable allows for asymmetric responses to different outcomes (*e.g*., wins vs. losses), and also captures the variable timing of each event in the task. The task variables were the following: (4) stimulus variables of left and right clicks and flashes (aligned to trial start). (4) Choice variables for choosing either the left of the right port (aligned to exit center port). Similarly, we included an alternate parameterization of choosing either the safe port (*p* = 1) or the risky port (*p* < 1). (5) Outcome variables were wins or losses on the current trial (aligned to choice); and reward history variables of previous wins, losses, or a previous opt-opt (aligned to trial start). We included a previous reward rate task variable that was calculated as the average rewarded volume from all previous trials in the session (aligned to trial start). We also included a “session progress” variable as the normalized [0,1] trial number in the session (aligned to trial start). This covariate captures motivational or satiety effects on firing rate over the course of a session. Finally, model comparison of cross-validated log-likelihood indicated that an autoregressive spike-history kernel was not necessary for our model.

#### 5.4.1 Parameter learning, hyperparameter choice, and model validation

The set of parameters θ of the model are the kernel weights 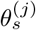 and the background firing rate *θ*_0_. *θ* were fit by minimizing 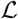, the negative-log likelihood – log[*p*(*y*|*θ*)] with an additional *L*_2_ penalty acting as a prior over model parameters [15]:

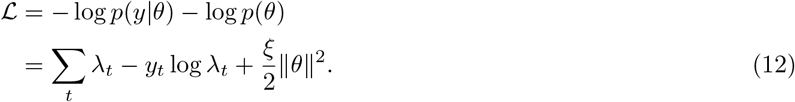

Model parameters were chosen through four-fold cross validation, and ξ was found through a grid search optimization on a held-out test set. Specifically, we split the data from each neuron into 5 equal parts that were balanced among trial contingencies (*i.e*., equal amounts of wins/losses, previous wins/losses, and left/right choices per partition). One partition (test set) was not utilized in fitting *θ*, and was held out for later model comparison and hyperparameter choice. The remaining four partitions were used in cross validation to fit four models, with each model fit on three partitions and assessed on the fourth “validation” partition. The model with the lowest negative log-likelihood on the validation set was chosen for further analysis. This procedure was repeated iteratively on an increasingly smaller grid of initial hyperparameter values ξ ∈ |10^−5^, 10], and the hyperparameter yielding the lowest negative log-likelihood on the test partition was chosen. We chose this approach for hyperparameter optimization in lieu of approximations such as calculating evidence [74], as we found the underlying approximations to be limiting for our data. In general, when building our model we assessed the aspects of other hyperparameter choices (*i.e*., kernel length, symmetric vs. asymmetric conditional kernels, number of task variables) with cross-validated negative log-likelihood on held-out test data. See the model comparison section of Supplementary Materials for further details. Finally, kernel covariances and standard deviations were estimated using the inverse of the Hessian of 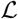.

#### 5.4.2 Basis functions

We utilized a log-scaled, raised cosine basis function set for our model [15]. These functions offer the ability to generate impulse-like responses shortly after stimulus presentation, as well as broader, longer time-scale effects. The form of the basis function is given as

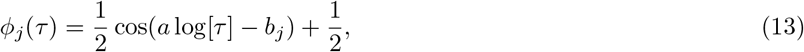

where *a* is parameter that controls breadth of support of each basis function, and *b_j_* controls the location of its maxima. The parameters *a, b_j_* were chosen to spread the set of basis functions {*ϕ_j_*} in roughly a log-linear placement across their range of support. This gives a better coverage than linear spacing, as the basis functions increase in breadth as *b_j_* increases. We used 9 such functions, and additionally augmented our basis set with two decaying exponential basis functions placed in the first two time bins to capture any impulse-like behavior at the onset of the task variable. This gives *M* = 11 basis functions for all kernels in the study. After a cursory model comparison of different kernel lengths we took a conservative approach and utilized a range of support of 4s for each kernel.

### 5.5 Coefficient of partial determination

The coefficient of partial determination (CPD) is a measure of the relative amount of % variance explained by a given covariate, and here we use it to quantify the encoding of behavioral information in single units. The CPD is defined as [30]

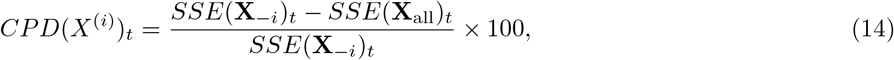

where *SSE* is the trial-summed, squared error between data and model. **X**_all_ refers to the full model with all covariates. **X**_–*i*_ implies a reduced model with the effect of *X*^(*i*)^ omitted. Since our model utilized a ont-hot encoding for each condition, we calculated CPD by omitting the following groups of covariates: 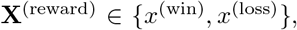, 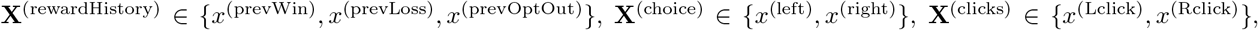, 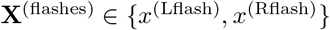.

Our measure of CPD assessed the encoding of the covariate of interest (*e.g*., previous win or loss) separately from encoding of the general event-aligned response (*e.g*., trial start). As such, our reduced model averaged the kernels that corresponded to the event-aligned response to create behaviorally irrelevant task variables. For example, for CPD of reward encoding: 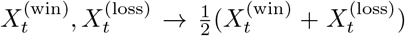. For reward, reward history, and choice CPD calculations, we additionally weighted each trial type in SSE such that each condition contributed equally to CPD, and omitted any bias due to an imbalance in trial statistics. Due to data limitations, we utilized the full set of training, testing, and validation data to calculate CPD. Units with a model fit of *R*^2^ > 0 (590/659 units) were used in CPD calculation.

We assessed the significance of the CPD result by comparing it to a null distribution of 500 CPD values that were generated by shuffling the trial labels among the relevant covariates (*e.g*., shuffle win and loss labels across trials, keeping timing of event and all other covariates fixed). CPD was deemed significant if it fell outside of the one-sided 95% confidence interval of the shuffle distribution, and plotted values in Figure 4 subtract off the mean of the shuffle distribution from the CPD.

### 5.6 Mutual information

Mutual information (MI) was used to calculate how much information about task variables is contained in lOFC firing rates, in different time windows throughout a trial. We calculated MI between spikes **Y_t_** and a group of covariates **X^(m)^** ∈ {*x_i_*} (detailed in the above section) as:

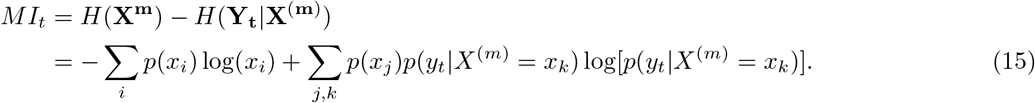

The first term is the entropy of the stimulus, which is calculated from the empirical distribution. The second term is the conditional entropy of spiking, and requires calculation of the conditional distribution *p*(*y_t_*|*X*^(*m*)^ = *x_k_*) by marginalizing over the fully conditional distribution that contains all other covariates from the model.

We modeled the fully conditional distribution *p*(*y_t_*|*X*^(1)^, *X*^(2)^, … *X*^(*S*)^, .. *X*^(*m*)^ = *x_k_*) via sampling of a doubly stochastic process, in which normal distributions of model parameters were propagated through our GLM and Poisson spiking. Specifically, for each time bin within each trial we sampled 500 parameter values from the normal distribution for each *θ*, where the covariance of *θ* was estimated as the inverse of the Hessian of 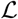. These samples were the passed through the exponential nonlinearity to generate a log-normal distribution of λ values, which were used as the rate parameter for a Poisson process that generated spikes. This spiking distribution of 500 samples was truncated at 10 spikes/50ms bin (a 200Hz cutoff), and the conditional distribution *p*(*y_t_*|*X*^(*m*)^ = *x_k_*) was then taken as the average over trials in which *X*^(*m*)^ = *x_k_*. Units with a model fit of *R*^2^ > 0 (590/659 units) were used in MI calculations. We assessed significance of MI in each time bin by comparing to an analogously created distribution of 500 MI values in which trial labels were shuffled. Significant MI fell outside of the 95% confidence interval of this distribution.

### 5.7 Waveform analysis

To investigate if different types of neurons may underlie different clusters, we extracted the action potential (AP) peak and after-hyperpolarization (AHP) peak from the waveform associated with each single unit. Waveforms were recorded on tetrodes, and the channel with the highest amplitude signal was used for waveform analysis. An initial manual curation of the waveforms excluded 18/659 units with poor isolation of the largest-amplitude channel, due to damage of the electrode. The remaining waveforms were standardized by upsampling the waveform by a factor of 20, and then mean-subtracting and normalizing by their maximum range. AP and AHP peaks were identified, then a threshold around zero (*α* = 0.05) was set to identify the beginning and end of the peak. We used the AP half-width rather than full width, as we found this to be a more robust measure of AP activity. Clustering of the two putative RSU and FSU units was performed using k-means.

### 5.8 Discriminability for reward history

The discriminability between previous wins and previous losses in Figure 5C was calculated in its unsigned form

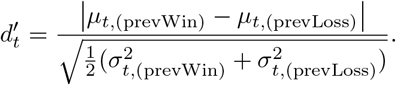

To account for an inflated range of nonsignificant *d*′ values around zero due to the unsigned form of *d*′, we subtracted from this quantity the mean *d*′ of a trial-shuffled distribution. The shuffled distribution was generated by shuffling trial labels 1000 times. Significant *d*′ values were identified as being outside of the one-sided 95% confidence interval of the shuffled distribution. Trial types were balanced when generating the shuffle distributions.

### 5.9 Linear regression models of pre-choice and post-choice epochs

For the linear regression of pre-choice epoch in Suppl. Fig. S11, the model used the trial history of wins and losses on the previous five trials as regressors, 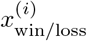, the choice on that trial, 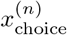, and an offset term, *c*_0_:

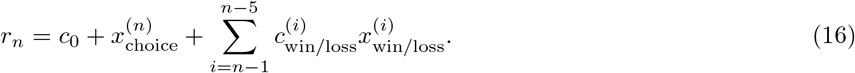

The regressors were binary +1/-1 variables for win(+1)/loss(−1) outcomes and left(+1)/right(−1) choice. Model coefficients c were fit with Matlab’s fitlm function, and significance was assessed via an F-test comparing to the baseline model containing only *c*_0_. The significance of each coefficient was assessed via a t-test with a cutoff of *p* < 0.05 for significance.

Similarly, the results in Figure 6 and Suppl. Fig. S14 investigating the adaptation of reward representations regressed *r_n_*, the average firing rate in the 1 second interval after reward onset on each trial *n* (post-choice epoch), against previous and current trial outcomes. The first model investigated adaptation of the rewarded outcome representation by including a binary win(+1)/loss(−1) regressor for current trial reward outcome, a binary win(+1)/loss(−1) regressor for previous reward outcome, the left(+1)/right(−1) choice on that trial, and an offset term :

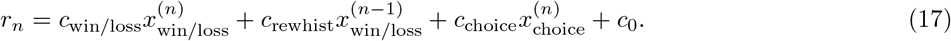

Similarly, the other model investigating adaptation of reward volume representations instead used a regressor for rewarded volume, and a binary loss(1/0) regressor for outcome on the current trial:

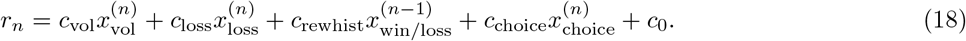

Finally, the regression in Suppl. Fig. S13 modeled the post-choice epoch using a reward prediction error as a regressor:

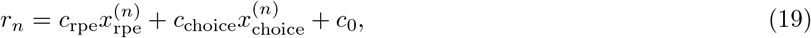

where the RPE regressor was the difference in rewarded volume and the expected value of the chosen option on that trial: 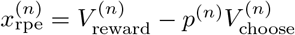.

## 6 Acknowledgements

We thank Carla Golden, Alex Piet, Abby Muthusamy, and Anne Churchland for helpful discussions and comments on the manuscript. This work was supported by NIMH grant 1R01MH125571-01 (CMC and CS).

## 7 Supplementary information

### 7.1 Excluded Data

The full data set consisted of 1881 single- and multi-units. Multi-unit recordings and neurons with a mean firing rate of less than 1 Hz were discarded, and the remaining 659 neurons were fit with the GLM, as well as utilized in the clustering procedures. Only neurons with a liberal threshold of *R*^2^ > 0 were utilized in CPD and mutual information calculations (590 units). Additionally, we found that CPD measures for neurons can sometimes be noisy, and of the 659 neurons fit by the model, we further excluded neurons as outliers in the cluster-averaged CPD calculation if they had a CPD value > 0.5. This excluded a further 10 neurons.

### 7.2 GLM model comparison

To choose the best-fit model to our data we performed model comparison on held-out testing data. For comparison we considered models with additional task variables such as current and previously rewarded volume; as well as reduced models that omitted the variables relating to previous opt out trials, the previous reward rate, and the session progress. We also considered a reduced model that omitted the reward history contribution entirely. In each case, we assessed the population level change in model performance via a Wilcoxon signed rank test (Suppl. Fig. S7). Our chosen model demonstrated a significantly lower population median in its negative log-likelihood than other models. We note that while our chosen model performed better than the reduced model at the population level, the median changes in neural responses were relatively slight (Suppl. Fig. S7 C, left panel). However, some neurons showed relatively strong improvement from introducing previousOptOut, previousRewardRate, and sessionProgress; which motivated us to keep them for the entire population (*i.e*., Suppl. Fig. S7 C, middle panel). Additionally, we investigated a symmetrized form of our model that used a binary (±1) encoding of task variables, as opposed to a one-hot encoding. This model was rejected at the population level via model comparison (not shown).

### 7.3 Supplementary Figures

**Figure S1:**
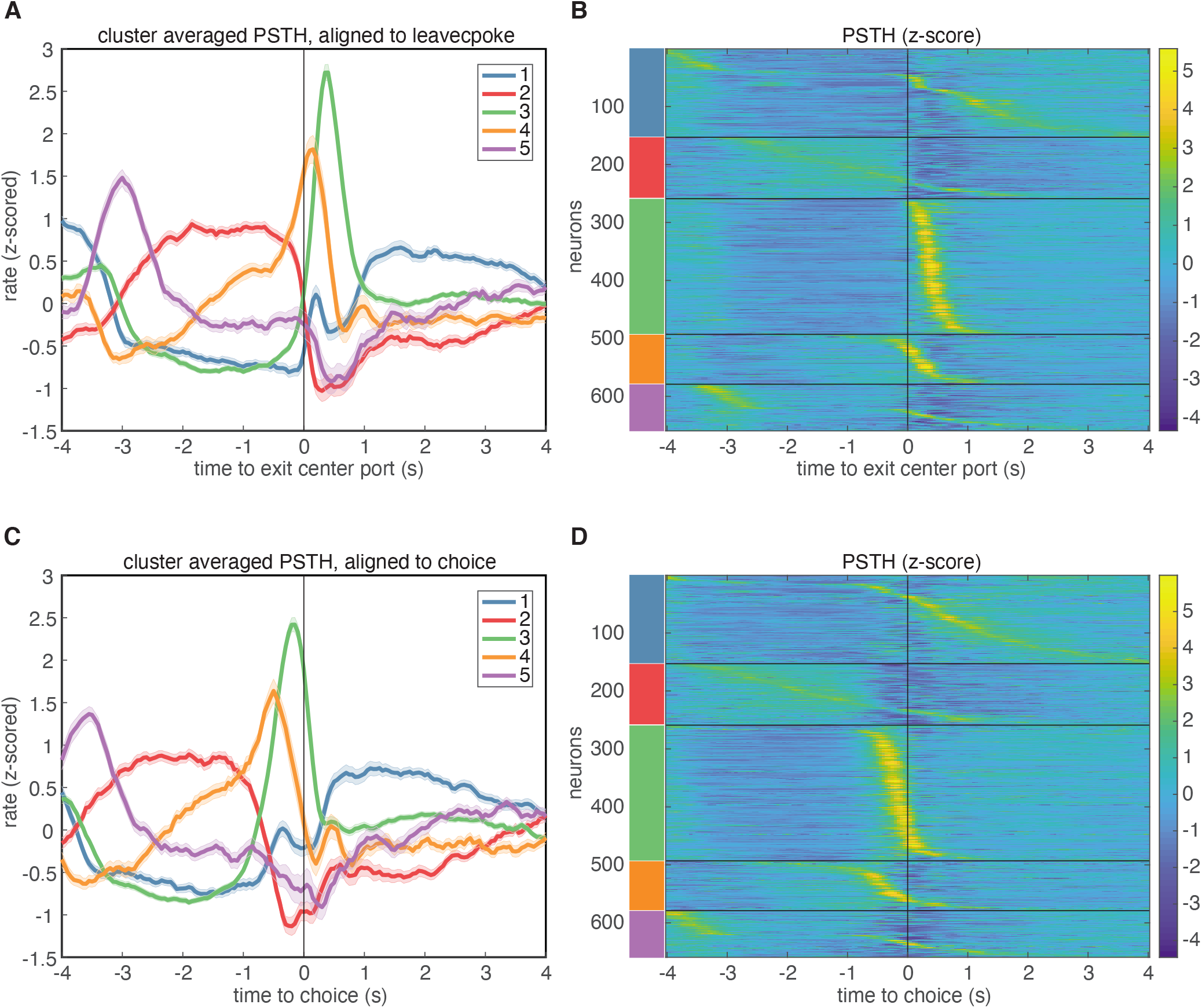
PSTH-based clustering of responses, aligned to different events in the task. A-B. Results aligned to exiting the center port. C-D. Results aligned to choice.

**Figure S2:**
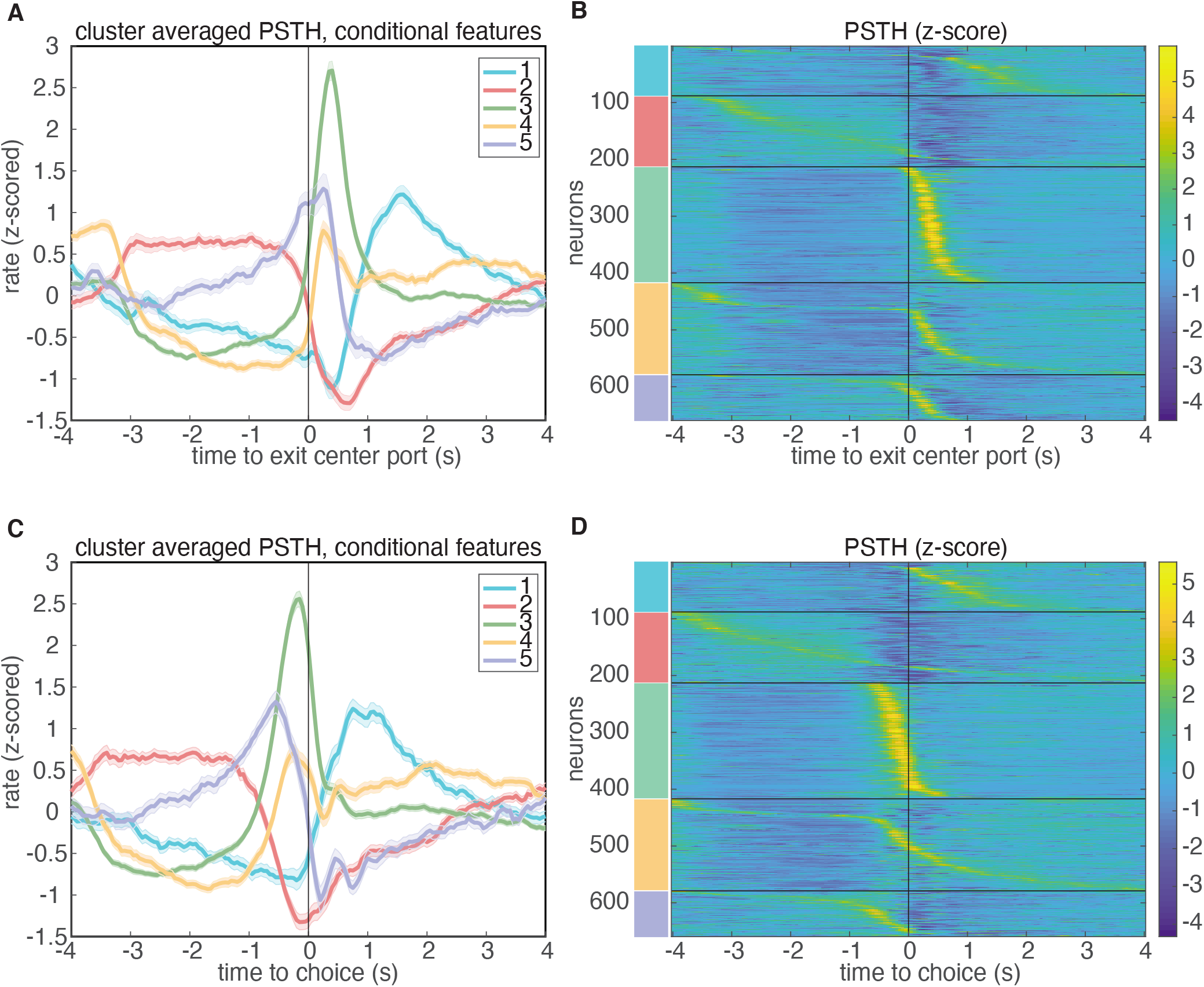
Results of clustering on conditional features space, aligned to different events in the task. A-B. Results aligned to exiting the center port. C-D. Results aligned to choice.

**Figure S3:**
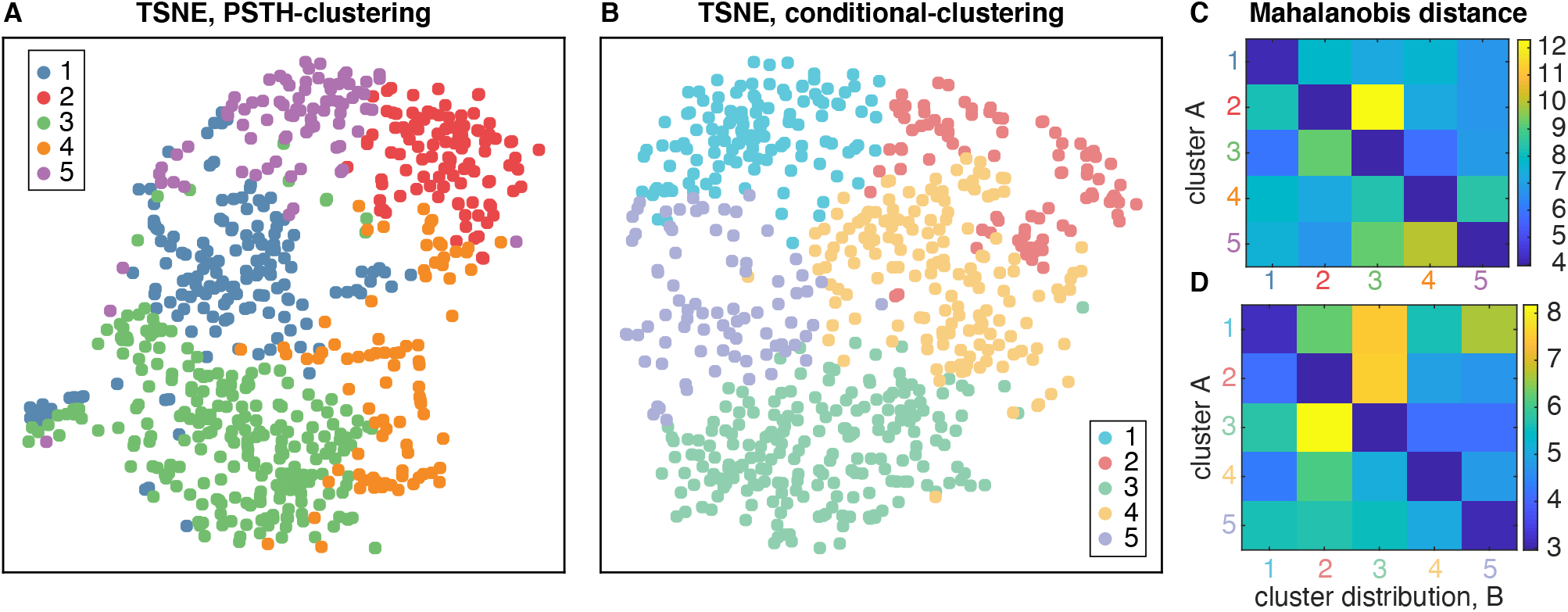
Similarity of clusters in feature space. A. t-SNE embedding of the PSTH feature space: Dots are individual neurons, and colors denote the cluster identity. B. Similar t-SNE embedding of the conditional feature space. Quantification of the cluster similarity through the cluster-averaged Mahalanobis distance of neurons from a given cluster distribution from all other clusters for C. the PSTH feature space clustering result and D. the conditional feature space clustering result.

**Figure S4:**
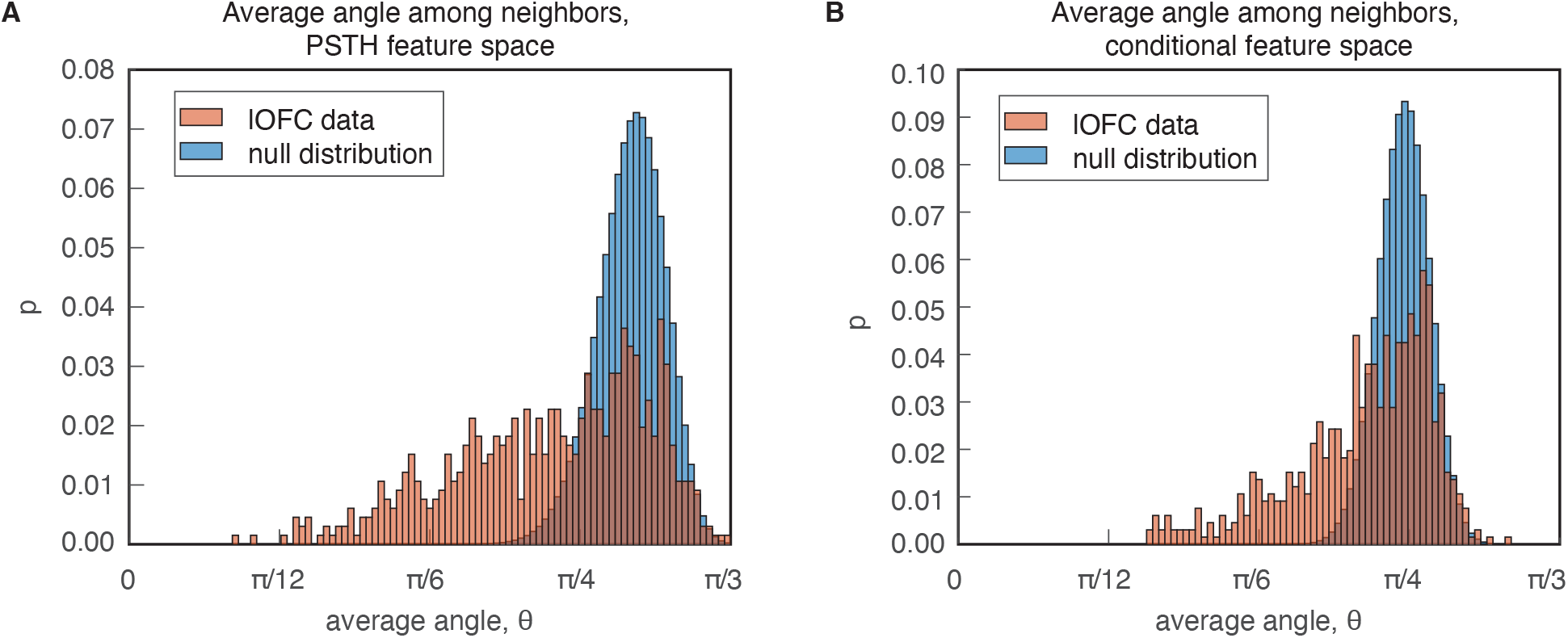
PAIRS statistic analysis. A) Average angles between k=3 nearest neighbors in the PSTH-based feature space (red) compared to a Gaussian distribution in the same dimension (blue). B) Similar distribution for the conditional feature space representation of neural responses, for k=8 nearest neighbors. The Kolomolgorov Smirnov (KS) statistic comparing the data distributions to the reference distribution yielded *p* ≈ 10^−125^ for the PSTH clustering, and *p* ≈ 10^−72^ for the conditional clustering. PAIRS statistics were 0.035 (*p* ≈ 10^−229^) and 0.06 (*p* ≈ 10^−59^). Details on the reference data are provided in Methods.

**Figure S5:**
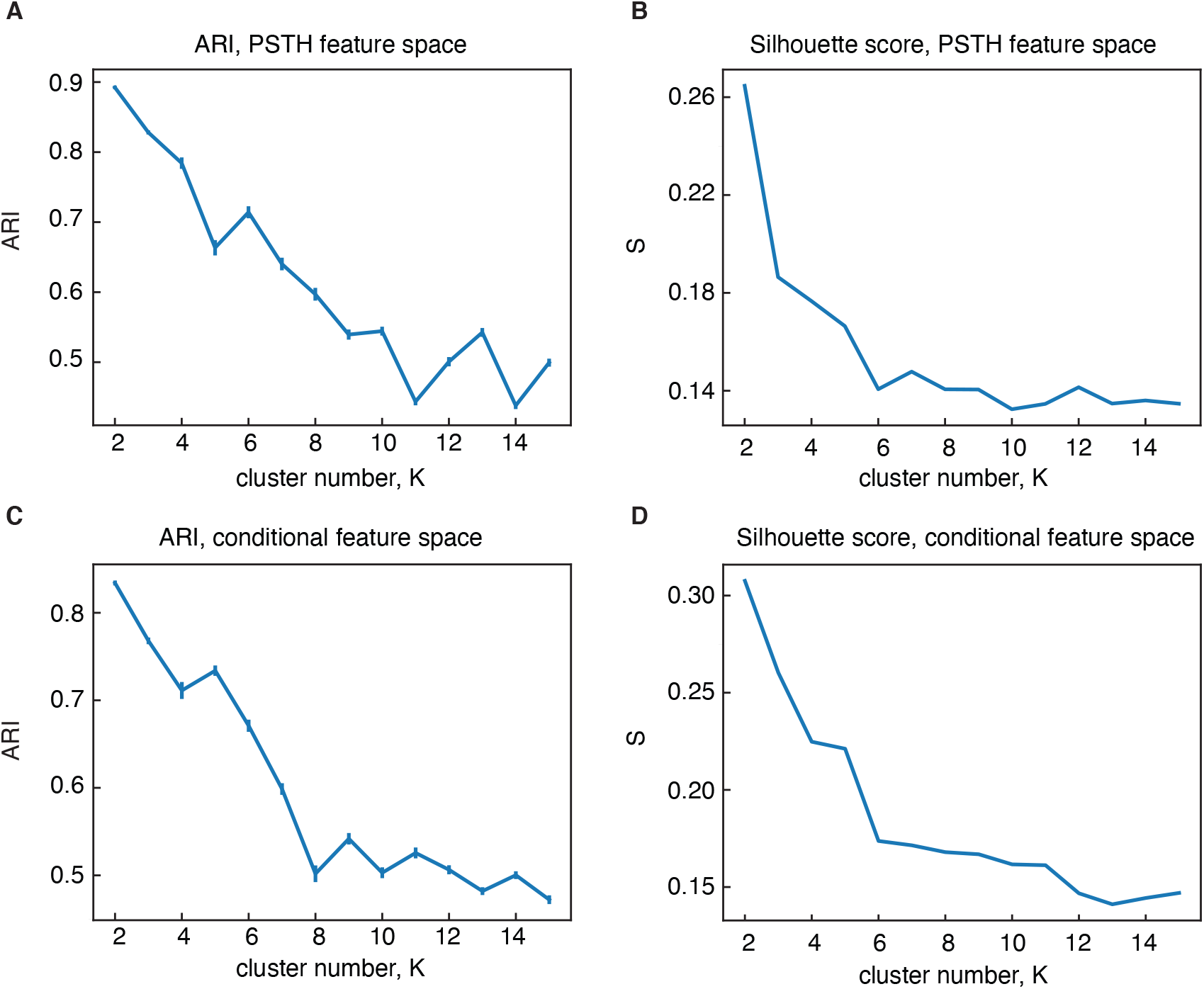
Silhouette score and adjusted Rand index (ARI) in different features spaces. A. Mean ARI values for clustering on PSTHs. ARI was calculated between the full dataset and 100 samples of subsampled data, created by sampling 90% of the population without replacement. Error bars are s.e.m. B. Silhouette score for clustering on PSTHs. C-D. Same analysis, but for the conditional feature space.

**Figure S6:**
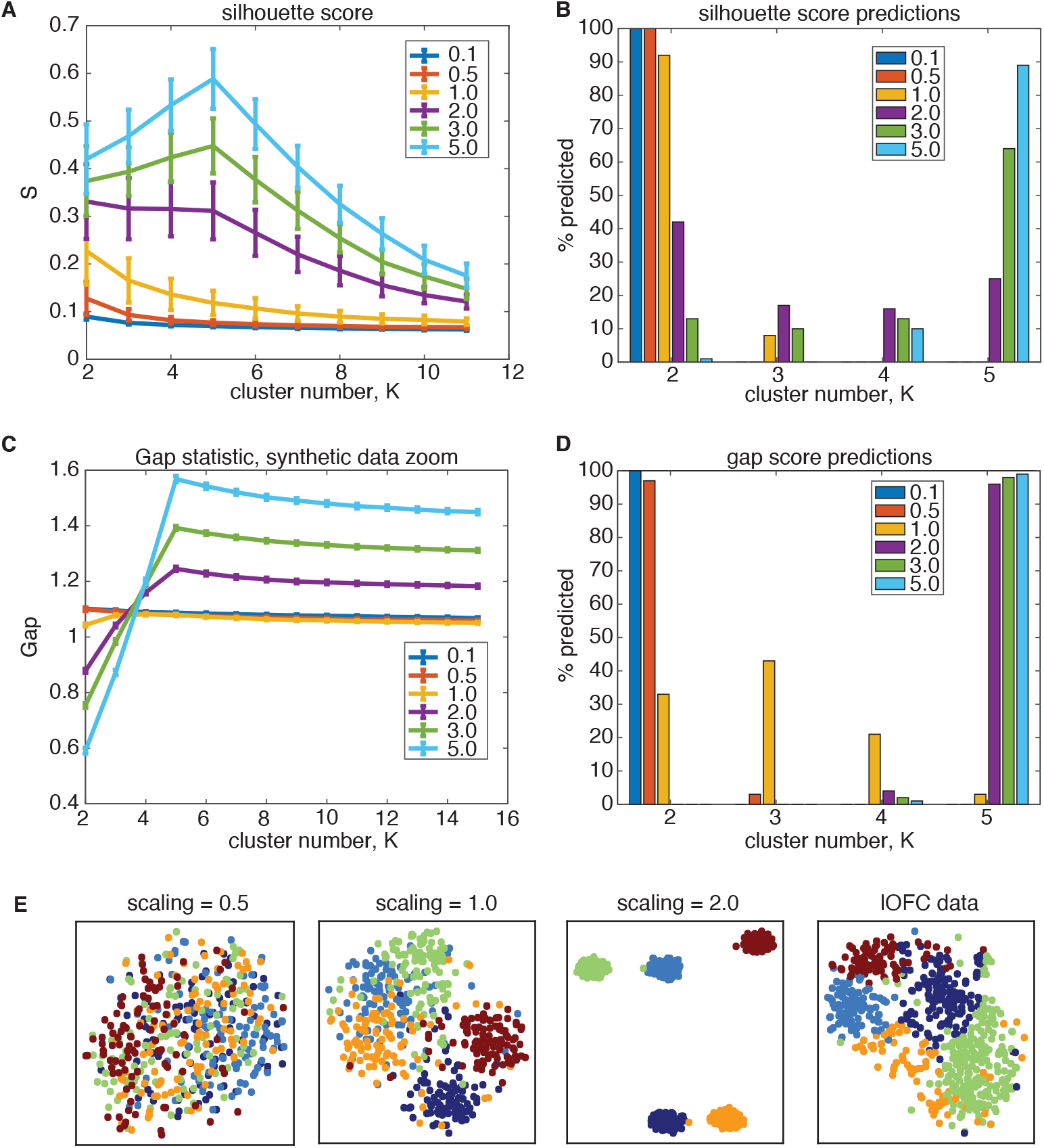
Silhouette score underestimate the number of clusters on ground-truth data in the same data regime as lOFC responses. The covariance structure lOFC data in the PSTH feature space was used to generated ground-truth, synthetic data containing *K* = 5 clusters, and the total-data covariance was scaled to investigate the ratio of across-cluster and within-cluster variance (See Methods for full details). A. Performance of silhouette score for varying scaling of total-data covariance, where scaling = 1.0 (yellow) is the same regime as lOFC responses. Error bars are 1 s.e.m over 100 ground-truth datasets B. Frequency of cluster number chosen for each dataset when using the silhouette score. For data scaled similarly to lOFC responses, silhouette scores predominantly choose K=2 clusters, and does not identify the true cluster number until the data is scaled to be very separated (scaling = 5.0, light blue) C. Performance of the gap statistic on the same synthetic data, where error bars are 1 s.e.m. D. Frequency of chosen cluster number using the gap statistic. The gap statistic also understimates cluster number in the lOFC data regime, but does so in a less severe manner: It predicts a range of values for scaling = 1.0, and quickly converges to predicting the correct number of clusters at only modest scaling of the data (scaling = 2.0, purple). E. t-SNE visualization of ground-truth data for different total-data covariance scaling.

**Figure S7:**
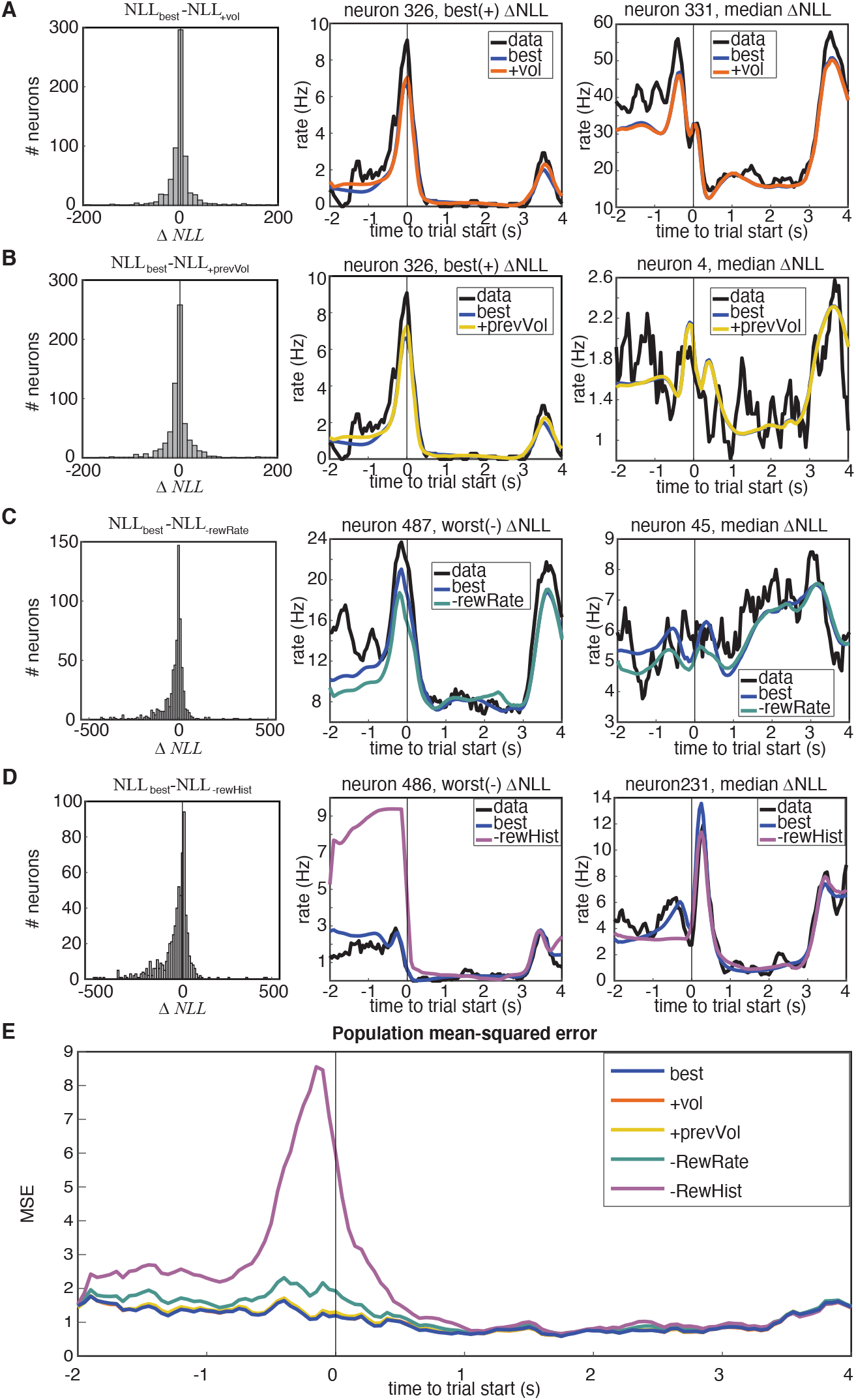
Model comparison. Model “best” was the model utilized in this work, and detailed in Figure 3. Two more complex models (+vol, +prevVol), as well as two simpler models (-RewRate,-RewHist) were compared through model comparison on held-out data log-likelihood. +prevVol substituted previous wins with rewarded volume on the previous trial, and +vol additionally substituted trial wins with rewarded volume. -RewRate omitted the session-Progress, prevRewardRate, and previousOptOut kernels, and -RewHist further omitted the remaining reward history kernels. A-D. Individual model comparisons. (Left) Histograms of the population-level changes in held-out negative log-likelihood demonstrate when a model performs significantly better if the median Δ*NLL* is different than 0. Significance is assessed through Wilcoxon signed rank test. *p* < 0.001 in all comparisons. Median Δ*NLL* values were Δ*NLL*_best,+vol_ = −0.62, Δ*NLL*_best, + prevVol_ = −1.77, Δ*NLL*_best,-RewRate_ = −4.14, Δ*NLL*_best,-RewHist_ = −15.13. Sample neuron fits demonstrating the most extreme change in Δ*NLL* are shown in the middle columns, and neurons demonstrating the median change in Δ*NLL* are shown in right columns. E. Performance of models assessed through the mean-squared error on PSTHs from held-out test data as compared to model predictions.

**Figure S8:**
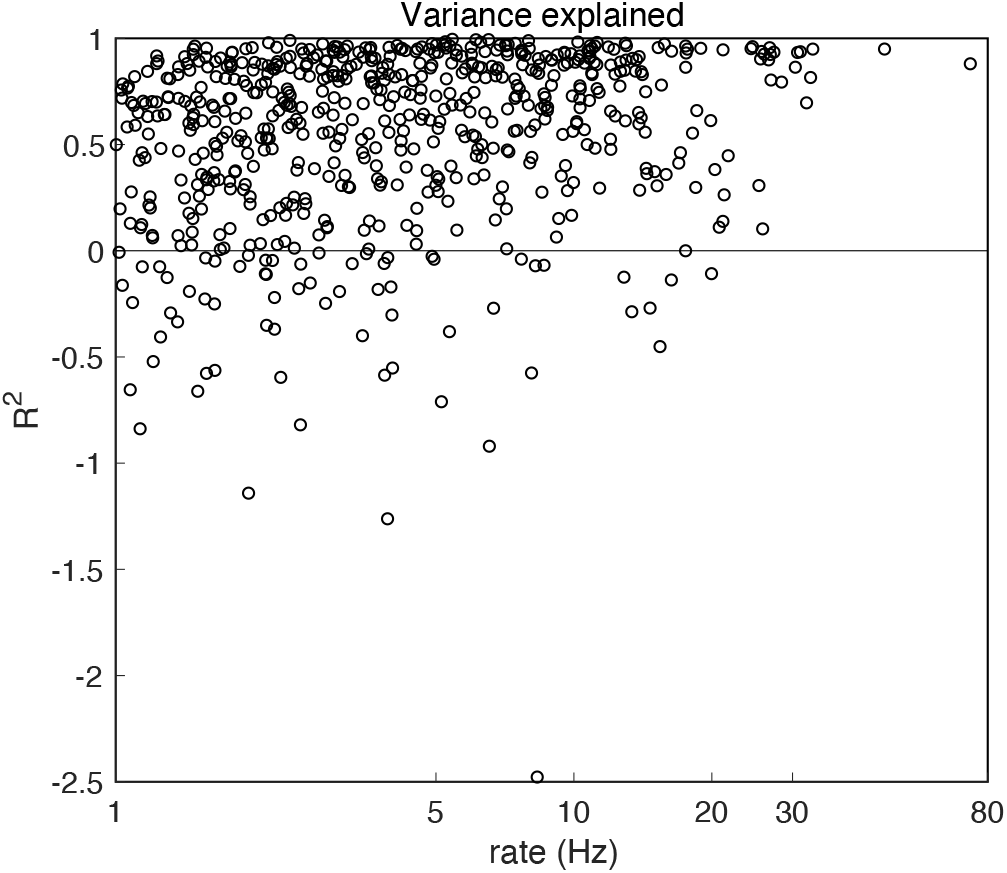
Variance explained, *R*^2^, for all model fits. 69 units (10.5%) of the models had *R*^2^ < 0, and were excluded from further analysis.

**Figure S9:**
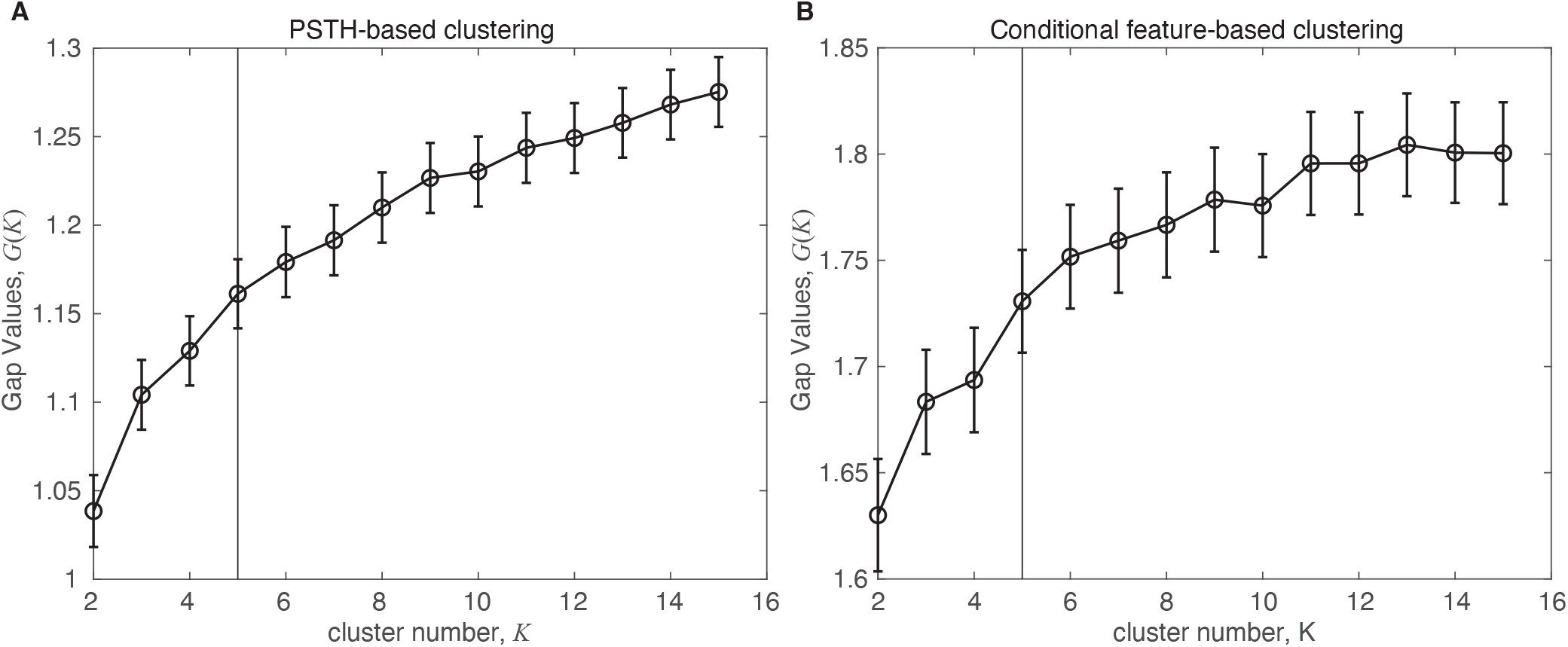
Gap statistic evaluation for both clustering approaches (see Methods for details). Error bars denote ± 2 s.e.m. A. PSTH-based clustering results. B. Alternative clustering approach based upon a conditional feature space. Vertical black lines denote largest significant number of clusters.

**Figure S10:**
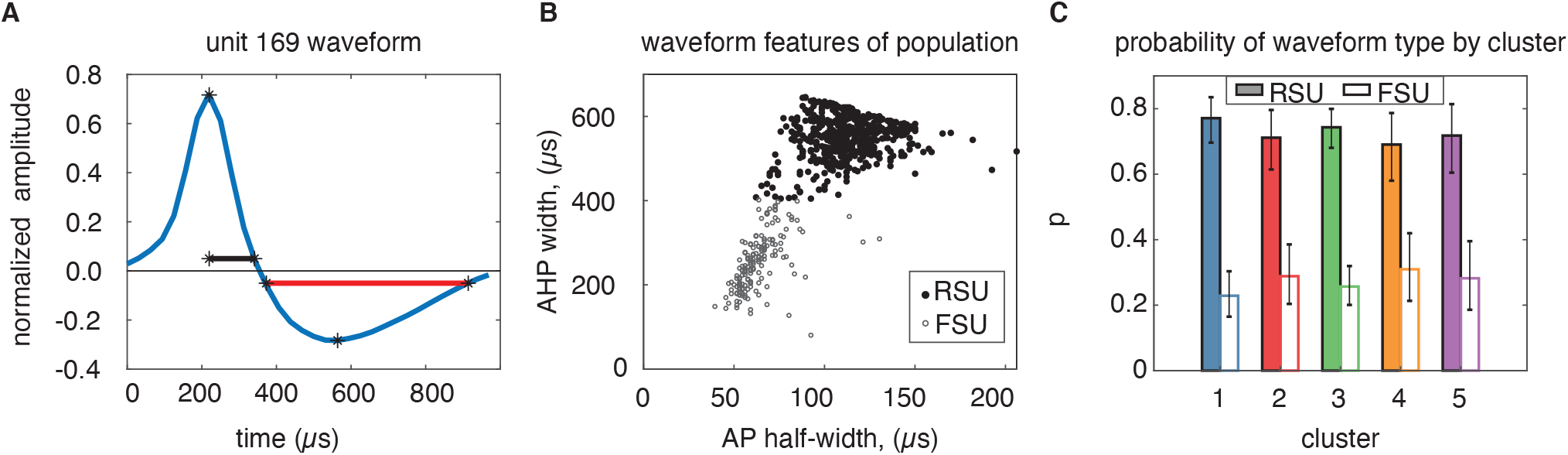
Waveform action potential (AP) and after-hyperpolarization (AHP) widths are equally distributed across clusters. A) the half-width of the AP peak (black), and full width of the AHP trough (red) were taken as key features of the waveform from each neuron. Waveforms were standardized by mean-subtracting and normalizing by the maximum amplitude range. B) AP and AHP for the population of units clustered into two putative groups: Regular spiking units (RSU, solid circles), and fast spiking units (FSU, open circles). C) Probability of an RSU or FSU neuron occurring in each N=5 clusters from the PSTH-based clustering labeling. Error bars denote the 95% binomial distribution confidence interval. The means of the two putative FSU and RSU clusters were significantly different (*μ_RSU_* = 4.65*Hz*, *μ*_FSU_ = 9.66*Hz*, *p* < 10^−4^, signed-rank test, not shown).

**Figure S11:**
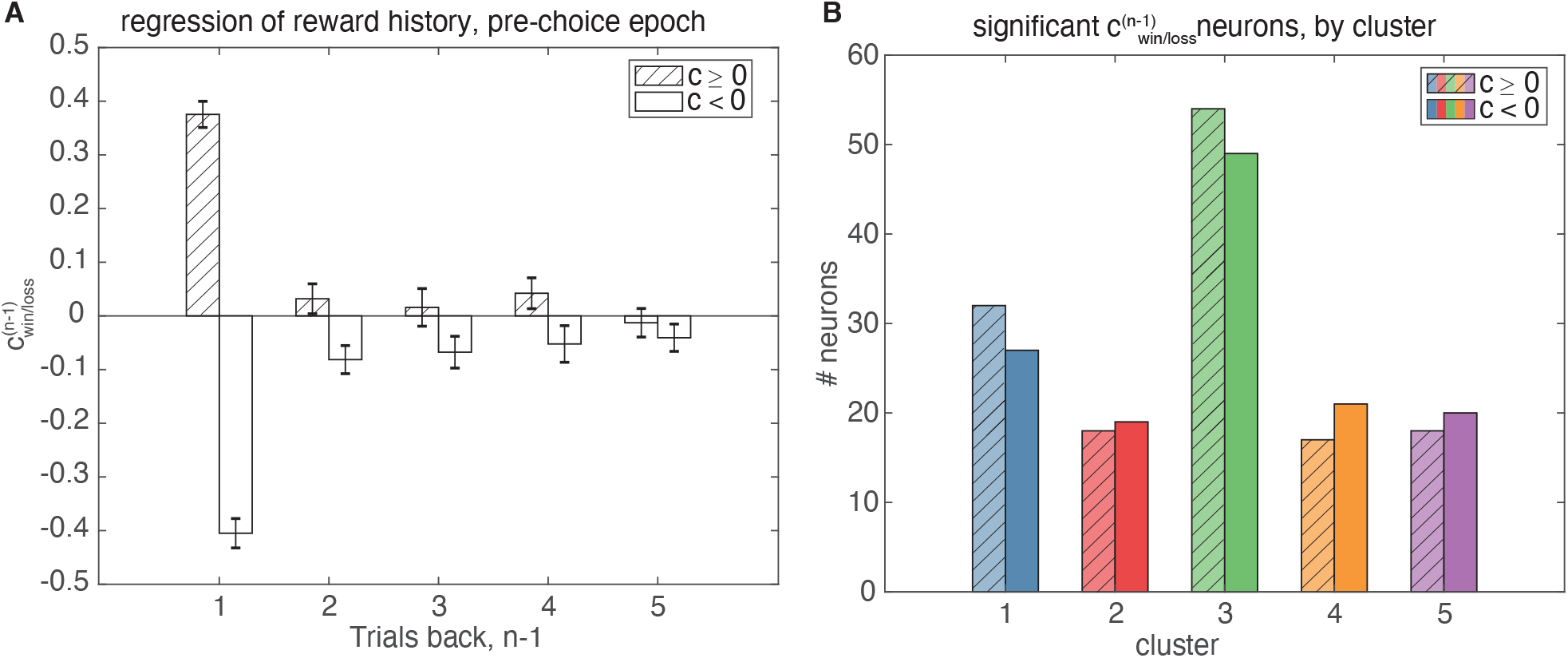
Linear regression of the pre-choice epoch based on recent trial history. A. Average regression coefficients for all statistically significant models with significant 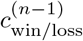 (*p* < 0.05 for F-test comparing full model to meanrate baseline model, *p* < 0.05 for t-test of 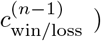). Regression coefficients one trial back have both positive and negative coefficients for reward outcome one trial back that convey different coding schemes for rewarded value, but coefficients for increasingly further trials back are not significant. B. Number of neurons with significant models containing either positive (shaded) or negative (not shaded) 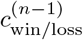 coefficients, separated by cluster. Cluster 3 contains the majority of statistically significant neurons.

**Figure S12:**
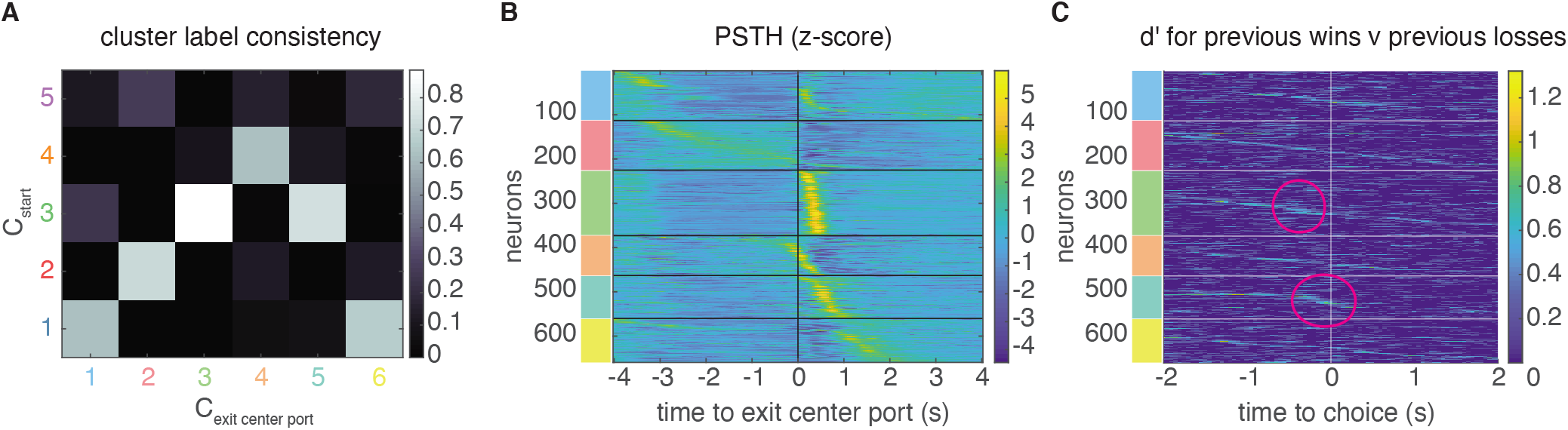
Clustering on PSTHs aligned to the later trial event of exiting the center port yields consistent responses to start-aligned PSTH clustering. A. Consistency of cluster labeling, calculated as the conditional probability *P*(*C*_exit center port_|*C*_start_) of belonging in any cluster from the exit center port clustering procedure (*C*_exit center port_), given that a unit belongs to a certain start-aligned PSTH cluster (*C*_start_). Responses in clusters 1-4 in *C*_start_ are similar to *C*_exit center port_, and cluster 5 in *C*_exit center port_ is also similar to cluster 3. B. Z-scored PSTH responses within each cluster, sorted by cluster and time to peak. Response profiles are similar to the original clustering procedure (compare to Fig. S1B). C. Sensitivity, *d*′, across time and neurons for reward history has more detailed structure when clustering on PSTHS aligned to later trial events. Cluster 3 still maintains sensitivity just before choice, though additional substructure is now visible in Cluster 5 right at choice on the current trial (magenta circles). *d*′ is sorted within the PSTH-based clusters by time-to-peak. Encoding of other task attributes, assessed through CPD was similar between the two clustering procedures (not shown).

**Figure S13:**
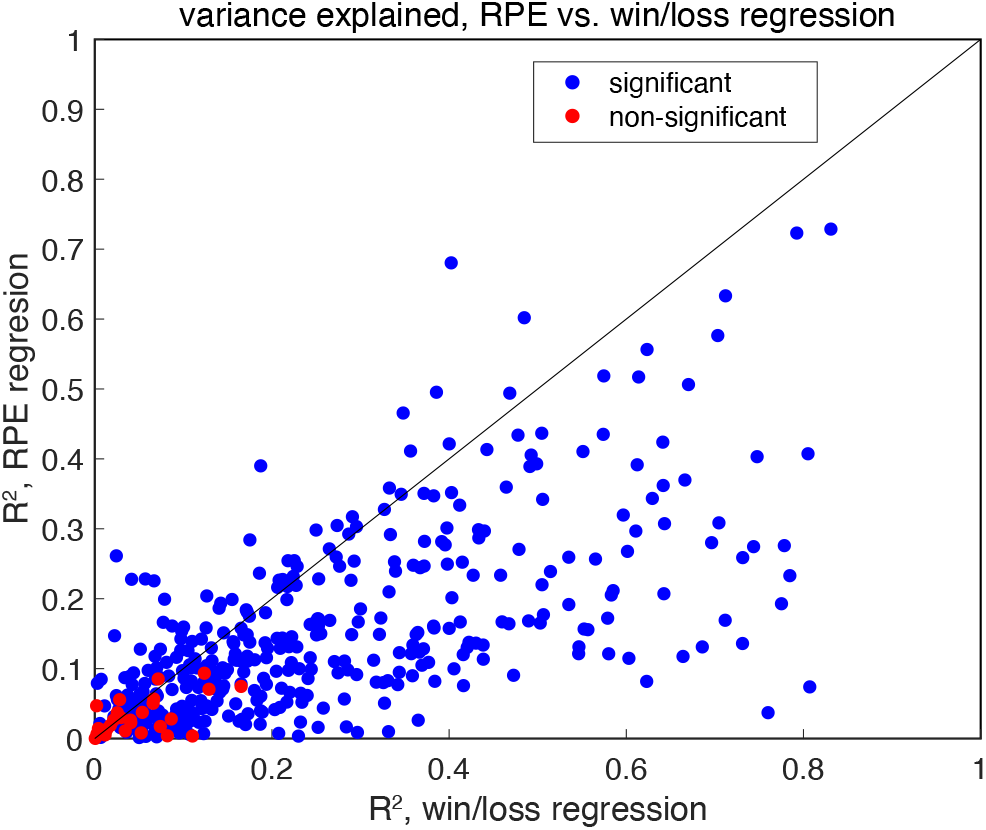
Linear regression of RPE in the post-choice epoch indicates that no RPE is present. Plotted is the variance explained for each neuron for two separate models: the binary win/loss model analyzed in Figure 6, and an equivalent model that replaces the current trial win/loss and past trial win/loss regressors with an RPE regressor. Blue dots indicate models with significant linear models, while red dots indicate non-significant models (threshold *p* < 0.05, F-test). The binary win/loss model captures more variance than the RPE model, indicating that wins and losses better explain that data. Further, model comparison between the two models by held out data log-likelihood reveals that the win/loss model is a better model (median Δ*NLL* = −5.9, *p* < 10^−4^, Wilcoxon signed rank test). The large proportion of RPE models that are significant is likely a reflection of the high correlation between RPE and win/loss regressors (*ρ* = 0.80).

**Figure S14:**
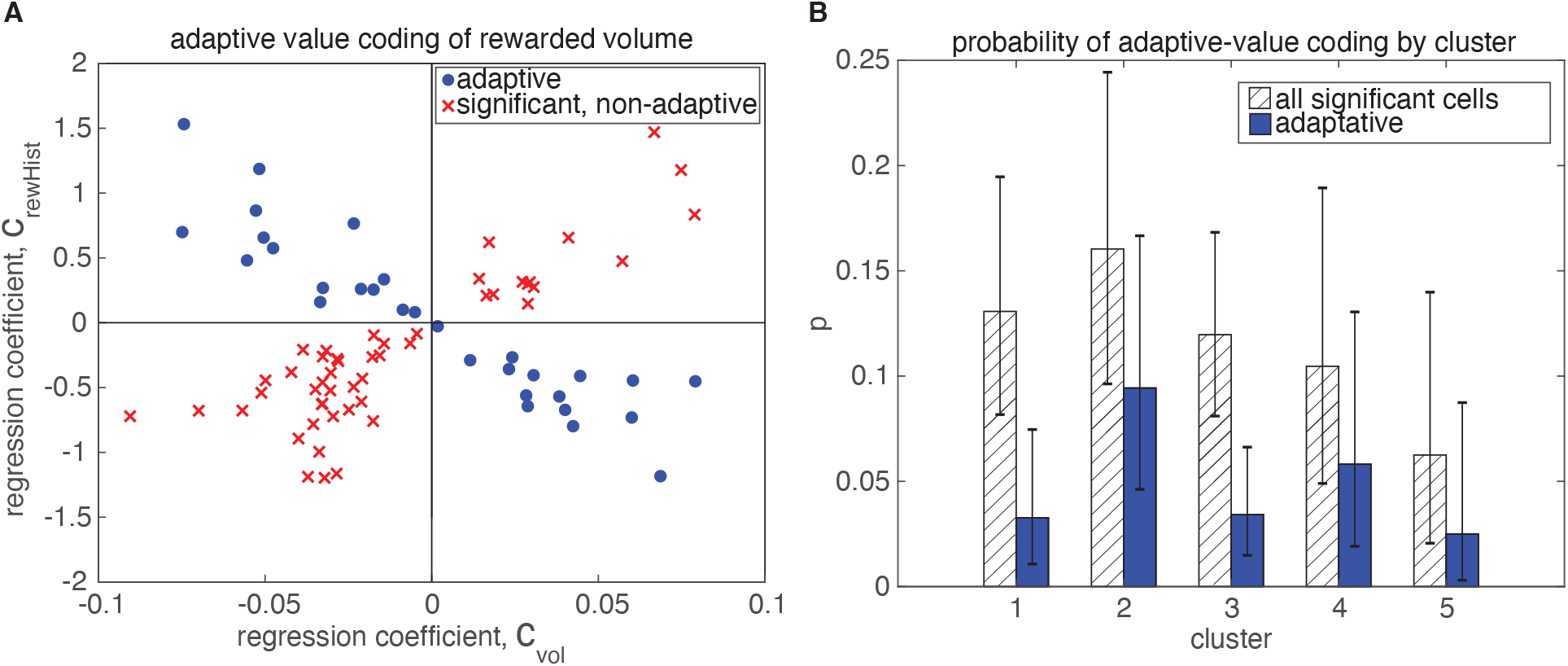
Adaptation of rewarded volume representation is present in fewer neurons than the reward outcome representation, but is still distributed across all clusters. Figure convention is similar to Fig. 6.

## Notes

### Competing Interest Statement

The authors have declared no competing interest.

